# *BIRC5* dependency defines a targetable vulnerability in *TP53* mutant acute myeloid leukemia

**DOI:** 10.1101/2025.05.17.654633

**Authors:** Ahmed M. Mamdouh, Fang Qi Lim, Yang Mi, Elyse A. Olesinski, Shaista Jasdanwala, Xiao Xian Lin, Edward Ayoub, Jordan Yong Ming Tan, Karanpreet Singh Bhatia, Yichen Wang, Shradha Garnaik, Kunal Tomar, Valeriia Sapozhnikova, Cheryl Gek Teng Chan, Chuqi Wang, Aarthi Nivasini Mahesh, Philipp Mertins, Brandon D. Brown, Torsten Haferlach, Melissa Ooi, Camille Lobry, Alexandre Puissant, Justin Taylor, Hussein Abbas, Steven M. Kornblau, Peter G. Miller, Sanjukta Das, Jan Krönke, Koji Itahana, Michael Andreeff, Shruti Bhatt

## Abstract

*TP53* mutations across multiple cancers, including acute myeloid leukemia (AML), are associated with poor outcomes irrespective of treatment modality. However, druggable vulnerabilities beyond canonical p53 targets remain largely unexplored. We identify *BIRC5* (encodes survivin), an inhibitor of the apoptosis protein (IAP) family, as a novel vulnerability in *TP53* mutant AML using an unbiased, comprehensive multiomics approach — whole-genome CRISPR knockout screen, bulk and single-cell RNA-seq, proteomics, and high-throughput drug screen. Mechanistically, *BIRC5* deletion in AML restored caspase-9 and -3/7 activity and downregulated other IAPs, implicating *BIRC5* as the central post-mitochondrial regulator for blocking apoptosis. p53 stabilization suppressed *BIRC5* selectively in *TP53* wild-type AML, explaining *BIRC5* upregulation in *TP53* mutant lines and AML primary tumors (n > 700). Longitudinal single-cell RNAseq (n = 22 pairs) revealed expansion of *BIRC5^high^* stem and progenitor leukemia clones in *TP53* mutant AML patients post-VenAza therapy. Survivin and IAP inhibitors emerged as top combination partners with VenAza in *TP53* mutant AML cells and showed potent *in vivo* leukemic blast inhibition in cell line and patient-derived xenograft models along with primary tumors. Beyond AML, *BIRC5* was upregulated broadly across 17 of 25 *TP53* mutant cancers in the TCGA cohort, and combination with survivin inhibitors overcame chemotherapy resistance in *TP53* deficient triple negative breast and colorectal cancers. These findings define *BIRC5* as a critical, targetable dependency and unveil survivin/IAP inhibition as a promising therapeutic axis to overcome p53-related resistance across both hematologic and solid malignancies.

**Key Points:**

1. BIRC5 upregulation is a novel dependency in TP53 mutant AML that mediates therapy resistance by evasion of apoptosis.
2. Combination with Survivin/IAP inhibitors overcomes venetoclax/azacitidine resistance in TP53 mutant AML.

## INTRODUCTION

*TP53* mutations or deletions are key drivers of therapy resistance in many cancers, resulting in poor responses to chemotherapy and targeted therapy.^1–3^ Among cancers harboring *TP53* mutations, acute myeloid leukemia (AML) is an aggressive hematological malignancy with especially dire outcomes. *TP53* mutations occur in approximately 10% of *de novo* AML, 20-35% of therapy-related myeloid neoplasia, and 70-80% of complex karyotype AML (CK-AML).^4,5,6^ The introduction of venetoclax (BCL-2 inhibitor) combined with azacitidine (hypomethylating agent) (VenAza) has transformed outcomes for many AML patients^7^, particularly in patients with poor-risk cytogenetics with wild-type *TP53* that historically responded poorly to 7+3 chemotherapy.^8–11^ However, this therapeutic benefit is blunted in patients with *TP53* mutations, where co-occurrence of poor-risk cytogenetics and *TP53* mutations results in drastically reduced median overall survival from 23.4 months to just 5.2 months.^7^ This emphasizes that *TP53* mutations alone are major contributors of VenAza resistance, even without poor-risk cytogenetics. Broadly, *TP53* mutant AML patients exhibiting significantly shorter relapse-free survival (4.3 months vs 18.9 months) and median overall survival (5.2 months vs 19.4 months) compared to those with wild-type *TP53*.^11–13^ These consistently poor outcomes, regardless of treatment modality, underscore the pressing unmet need to develop therapeutic strategies that overcome therapy resistance in *TP53* mutant AML.

Despite decades of research establishing *TP53* as the master regulator of apoptosis, the precise mechanisms by which *TP53* mutations confer treatment resistance beyond dysregulation of BCL-2 family proteins remain incompletely understood. p53 triggers mitochondrial outer membrane permeabilization (MOMP) to induce apoptosis by transcriptionally activating pro-apoptotic proteins, BAX, PUMA, and NOXA^14,15^, or by repressing anti-apoptotic proteins, MCL-1, BCL-2 and BCL-xL.^16,17^ Paradoxically, although MOMP is considered the ’point-of-no-return’ in apoptotic signaling across diverse cell types undergoing therapeutic stress, evidence suggests that cells expressing wild-type p53 may sometimes recover following MOMP activation.^18^ Alternatively, cytoplasmic p53 can be sequestered by anti-apoptotic BCL-2 family proteins, with disruption of these complexes by PUMA releasing p53 to trigger MOMP and apoptosis.^19–21^ Given the intricate interplay between p53 and BCL-2 family proteins, we hypothesize that other unexplored functions of wild-type p53 must exist that become dysregulated to promote apoptosis evasion in the mutant settings.

In this study, we utilized a systematic, unbiased multiomics approach to identify vulnerabilities for *TP53* mutant cells. We discovered that *TP53* mutant AML is functionally dependent on *BIRC5* (encodes survivin), an inhibitor of apoptosis protein (IAP), to mediate therapy resistance via inhibition of caspase-3 activation. We established that wild-type *TP53* negatively regulated *BIRC5* expression, providing a mechanistic explanation for *BIRC5* upregulation in *TP53* mutant/deficient cells. These findings reveal a previously unrecognized p53-dependent regulation of post-mitochondrial apoptotic signaling, the dysfunction of which drives therapy resistance. In our high-throughput screen, survivin/IAP inhibitors scored highly and demonstrated efficacious anti-leukemic effects in combination with VenAza *in vivo*. Crucially, this targetable *BIRC5* dependency represents an exploitable vulnerability in *TP53* mutant/deficient cancers, offering a novel strategy to overcome chemoresistance in these aggressive malignancies.

## MATERIALS AND METHODS

A detailed description of experimental methods can be found in the Supplemental Methods, including animal studies, caspase luminescence assay, cell culture, cell cycle, apoptosis, and viability studies, CRISPR library screen, cytotoxicity assay, data-independent acquisition proteomics, drug library screen, genetic editing in cell lines, reverse phase protein array, patient cohorts, RT-qPCR and RNA-seq, single cell RNA-seq, TMRE staining, and western blot.

### Statistical significance

Statistical significance was assessed using two-tailed t-tests unless otherwise indicated. Proteomics data were analyzed using two-sided t-test p-values adjusted by the Benjamini-Hochberg method to control for false discovery rate. One-tailed Wilcoxon rank-sum tests were used to compare the Munich Leukemia Laboratory, Beat AML, and TCGA cohorts and to evaluate differential survivin protein expression between *TP53* mutant versus wild-type and complex versus non-complex karyotype clinical samples. Differences in cumulative survival probabilities were determined using Kaplan-Meier analysis with long-rank testing. *In vivo* studies were analyzed using one-way ANOVA with Tukey adjustment. Statistical significance levels are indicated in all figures as follows: *p < 0.05, **p < 0.01, ***p < 0.001, ****p < 0.0001.

## RESULTS

### *BIRC5* is a dependency in *TP53* mutant/deficient AML for therapy resistance

Despite the widespread clinical use of VenAza in AML, resistance remains a major challenge in the *TP53* mutant setting. Hence, we sought to identify actionable dependencies that promote sensitivity to VenAza in *TP53* mutant and deficient AML. We conducted genome-wide CRISPR knockout screens in isogenic MOLM-13 AML cells (wild-type *TP53* (*TP53^+/+^*), *R248Q* missense mutation (*TP53^-/R^*^248^*^Q^*), *TP53* deficient (*TP53^-/-^*)) treated with either DMSO or VenAza (Figure 1A). The *R248Q* mutant was selected because it is the second most common *TP53* mutation in myeloid tumors, has poor prognosis, and conferred highest resistance to VenAza.^22–25^ CRISPR screens were performed using the Brunello library (77,441 sgRNAs targeting 19,114 genes).^26^ Functional dependencies in a single *TP53^+/+^* clone closely mirrored those of the parental MOLM-13 cell line (Figure S1A). In *TP53^-/R^*^248^*^Q^*and *TP53^-/-^* cells, we identified sensitizer or resister candidates at baseline (n = 35) and after VenAza (n = 262) (log2 fold change >1; Figure 1B). Within each genotype, comparison between DMSO and VenAza consistently showed enrichment in pro-apoptotic regulators (*BCL2L11, PMAIP1,* and *BAX*) yet no shared resistance genes across the top 100 gene hits (Figures 1C and S1B). This suggests that despite *TP53* mutations, apoptotic activators remain top genes to resensitize these tumors to VenAza. Beyond sensitizers, anti-apoptotic BCL-2 family proteins, *BCL2L1* and *MCL1*, were depleted in *TP53^-/R248Q^*cells following VenAza (Figure 1C).

**Figure 1.**
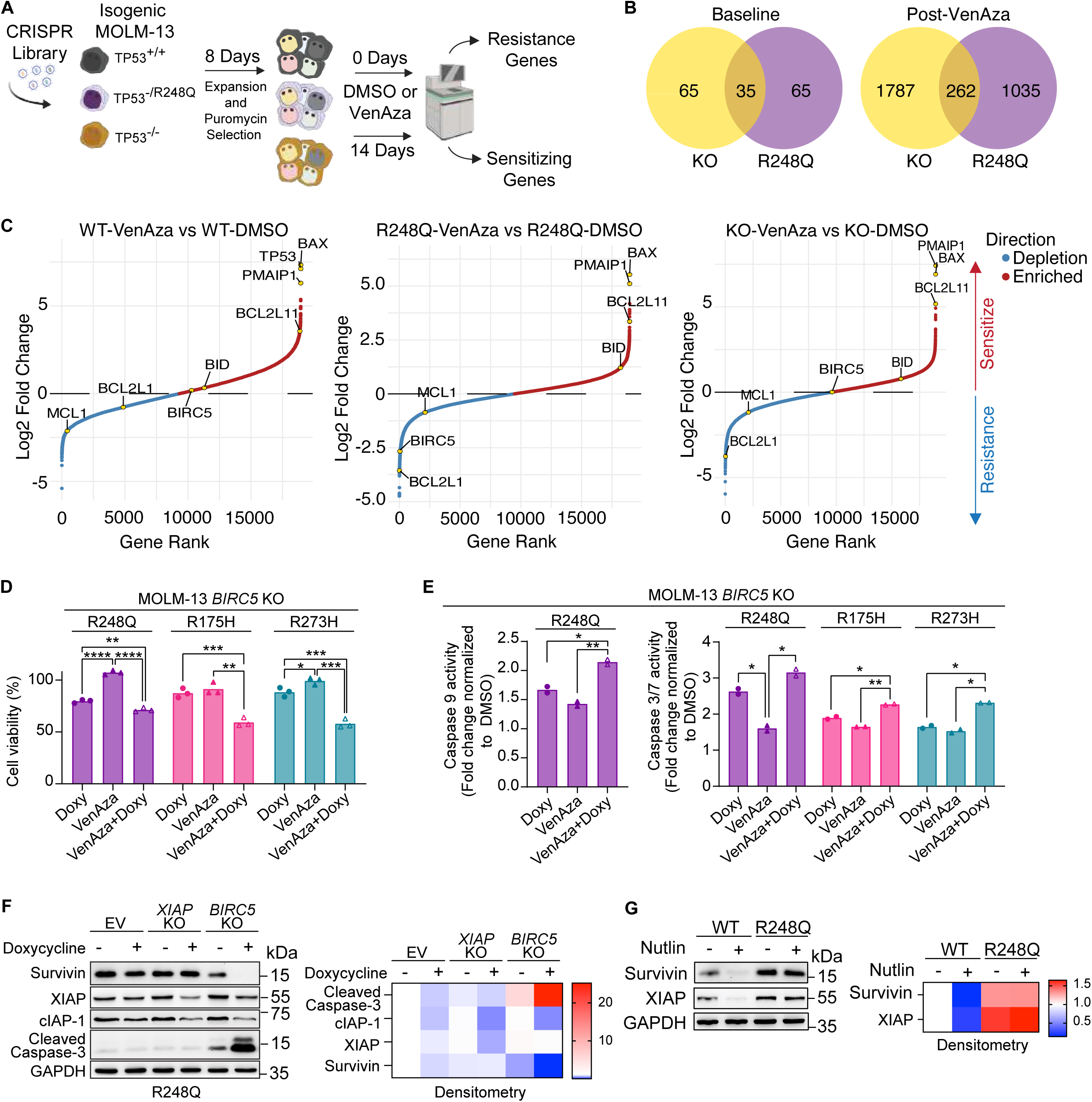
Unbiased CRISPR screen identifies *BIRC5* as a top resistance gene in *TP53* mutant AML. (A) Schematic of CRISPR screen for identifying sensitizing and resistance genes. Treatment with venetoclax (35 nM) and azacitidine (50 nM) for 14 days. Cells were sub-cultured in new media with treatment every three days. **(B)** Venn diagram of shared genes at baseline (35 genes) and following VenAza treatment (262 genes) across *TP53*^-/R248Q^ and *TP53^-/-^* cells. **(C)** Ranking plot of CRISPR screen for *TP53*^+/+^, *TP53*^-/R248Q^, and *TP53^-/-^* MOLM-13 cells treated with VenAza and DMSO. “Enriched” refers to genes that conferred higher killing (enriched sgRNA guides) and “depletion” refers to genes that conferred survival (depleted sgRNA guides) in response to VenAza. Cut-off was pre-determined by MAGeCKFlute, where cut-off scores for *TP53*^+/+^, *TP53*^-/R248Q^, and *TP53^-/-^* cells were ±2.466, ±1.328, and ±1.738, respectively. **(D-E)** *TP53*^-/R248Q^, *TP53*^-/R175H^, and *TP53*^-/273H^ cells with *BIRC5*-KO treated with doxycycline (10 ng), VenAza, or combination for 72 hours: **(D)** cell viability and **(E)** caspase-3/7 and caspase-9 assays. All data was normalized to DMSO (non-doxycycline)-treated cells. Analysis via ANOVA followed by Tukey for multiple test corrections. **(F)** Western blot and densitometry for indicated proteins with *BIRC5*-KO (+doxycycline) and *XIAP*-KO (+doxycycline). **(G)** Western blot and densitometry show that nutlin-3a (5 μM for 24 hours) downregulates survivin and XIAP selectively in MOLM-13 *TP53*^+/+^ cells but not *TP53*^-/R248Q^ cells. For all figures unless otherwise specified, treatment with VenAza: venetoclax 3 nM, azacitidine 50 nM. ns = no significance, *p<0.05, **p<0.01, ***p<0.001, ****p<0.0001.

Notably, the inhibitor of apoptosis protein (IAP) family gene, *BIRC5* (encodes survivin), was significantly depleted in *TP53^-/R248Q^*cells. IAPs negatively regulate programmed apoptosis by inhibiting effector and executioner caspases.^27,28^ RT-qPCR analysis confirmed that *BIRC5* was significantly upregulated in *TP53^-/R248Q^*and *TP53^-/-^* cells at baseline and following VenAza (Figure S1C). In contrast, other IAP family members – *BIRC2* (encodes cIAP-1), *BIRC6* (encodes BRUCE), and *XIAP* – were variably expressed, underscoring *BIRC5* as a dominantly elevated IAP gene (Figure S1C). This prompted us to test *BIRC5* as a novel mediator of VenAza resistance in *TP53* mutant AML. Loss of *BIRC5* using doxycycline-inducible knockouts (*BIRC5*-KO) restored sensitivity to VenAza in *TP53^-/R248Q^*, *TP53^-/R175H^*, and *TP53^-/R273H^* cells (Figure 1D). Combining *BIRC5-*KO with higher doses of azacitidine and venetoclax led to increased cytotoxicity, though the effect was greater from venetoclax (∼10% and ∼30%, respectively) (Figure S1D). *BIRC5*-KO also increased caspase-9 and caspase-3/7 activity in *TP53^-/mut^* and *TP53^+/+^*cells, confirming that *BIRC5* mechanistically suppresses downstream caspase activation to drive apoptotic resistance (Figures 1E and S1E).

Survivin (encoded by *BIRC5*) promotes XIAP stabilization by antagonizing its RING E3 ubiquitin ligase domain.^29^ This prompted us to investigate the involvement of other IAP family members. In *BIRC5*-KO, XIAP and c-IAP-1 protein levels were markedly reduced, suggesting that survivin supports their stability (Figures 1F). Despite priors studies showing that XIAP directly inhibits caspase-9 and caspase-3/7^30,31^, *XIAP* deletion (*XIAP*-KO) in *TP53^-/R248Q^* cells did not restore caspase activity nor sensitize cells to VenAza-induced cell death (Figure S1F). These findings support a model in which *BIRC5*, rather than *XIAP*, serves as the primary indirect mediator of caspase inhibition, with other IAPs potentially acting in concert to reinforce apoptotic resistance in *TP53* mutant AML.

### Wild-type *TP53* suppresses *BIRC5*

To explain the dependency of *TP53^-/R248Q^* and *TP53^-/-^* cells on *BIRC5*, we hypothesized that wild-type p53 suppresses survivin and XIAP. Pharmacologic stabilization of wild-type p53 with nutlin-3a decreased expression of XIAP and survivin in *TP53^+/+^*cells, an effect lost in *TP53^-/R248Q^* cells and consistent with p53’s transcriptional repression of IAP family members (Figure 1G). We found no evidence of direct p53 binding to *XIAP* and *BIRC5* promoters in a published chromatin-immunoprecipitation (ChIP) dataset.^24^ This suggests that regulation occurs indirectly through p53-dependent transcriptional targets rather than direct promoter occupancy, as opposed to prior reports^32^ (Figure S1G). *TP53* mutations also frequently co-occur with 17p deletions, and *BIRC5* is located on 17q. In AML and six cancers with substantial 17p loss from The Cancer Genome Atlas (TCGA) cohort, *BIRC5* upregulation occurred alongside 17q gain, suggesting that genomic instability may contribute to tissue-specific *BIRC5* elevation (Figure S2A-B). These findings demonstrate that wild-type *TP53* negatively regulates IAPs to promote apoptosis, while *TP53* mutations relieve this repression to enforce a post-mitochondrial blockade of caspase activation and cell death.

### Survivin and IAPs are upregulated in *TP53* mutant/deficient AML isogenic cells

We performed global proteomic analysis of *TP53* mutant AML cells using data-independent acquisition mass spectrometry. Unsupervised clustering at baseline showed clear segregation based on *TP53* status (*TP53^+/+^*, *TP53^-/R248Q^*, or *TP53^-/-^*) (Figure S3A). Among pro-apoptotic BH3-only proteins, BAX and PUMA (encoded by *BBC3*) were significantly downregulated in *TP53^-/R248Q^* and *TP53^-/-^* cells compared to *TP53^+/+^* cells (Figure S3B). We next examined post-mitochondrial apoptosis regulators and observed that survivin and XIAP were consistently upregulated, while BRUCE was downregulated (Figure 2A). Although BRUCE loss typically disrupts caspase-9 and SMAC degradation^33^, immunoblotting revealed no consistent changes in these pathways (Figure 2B). At baseline, survivin was elevated in 3/6 *TP53^-/mut^* cell lines and XIAP in 5/6 lines. cIAP-1 was generally unchanged except in *TP53^-/R248Q^* cells, where it was increased. *TP53^-/-^* cells exhibited increased expression of survivin, XIAP, and cIAP-1. This pattern was validated in an additional *TP53^-/-^* MOLM-13 cell line (Figure 2C). Survivin was also increased in the *TP53^-/-^* MV-4-11 cell line. Collectively, these results demonstrate that *TP53* mutant AML cells sustain elevated levels of IAP family proteins.

**Figure 2.**
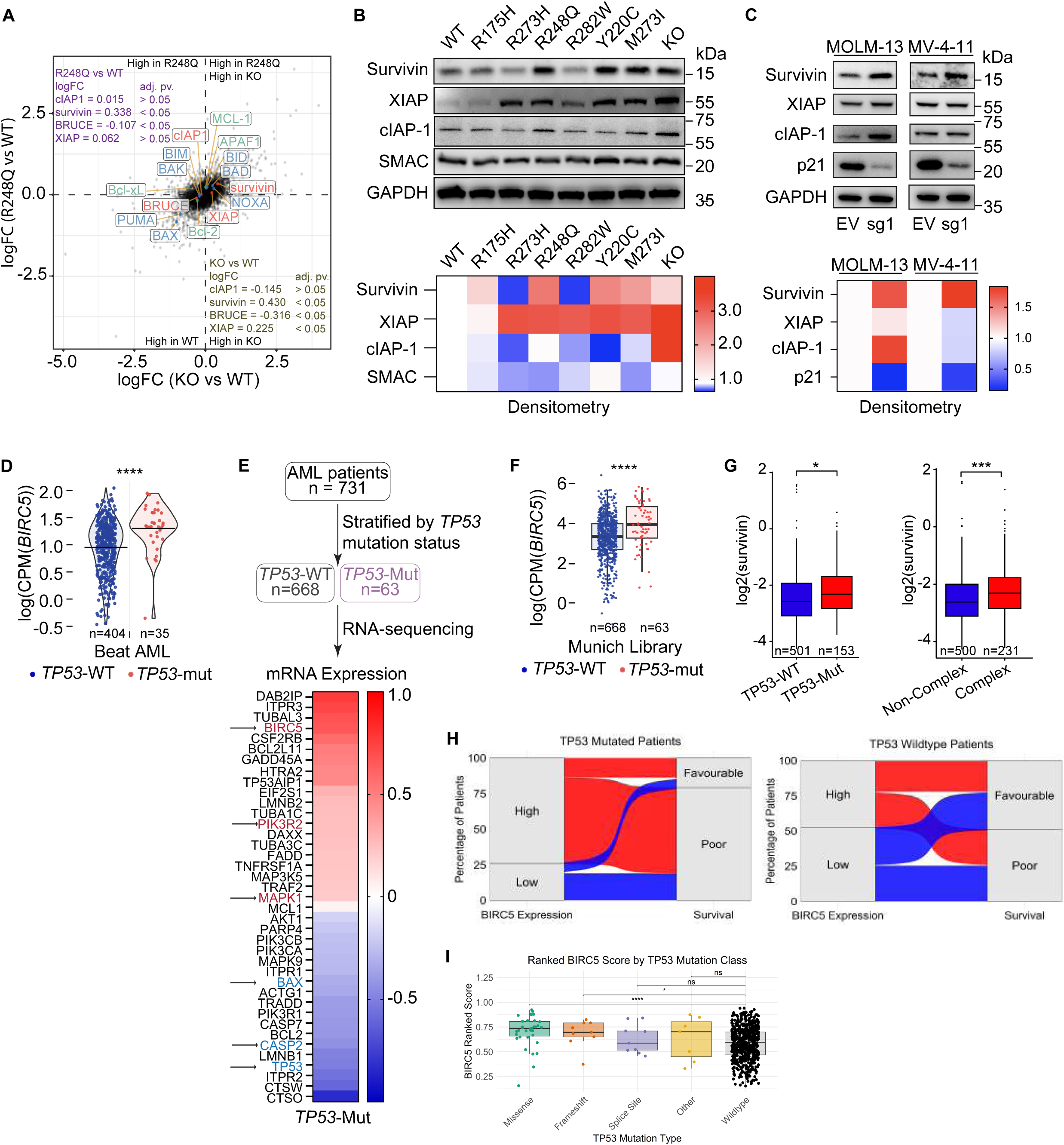
Primary AML tumors and isogenic cells with *TP53* mutations have upregulated *BIRC5* gene and protein expression. **(A)** Scatterplot of mass spectrometry in data-independent acquisition proteomics of isogenic *TP53*^-/R248Q^ and *TP53^-/-^* MOLM-13 cells compared to *TP53*^+/+^ cells at baseline. Blue: pro-apoptotic proteins; green: anti-apoptotic proteins; red: inhibitor of apoptosis proteins. Analysis via two-sided t-test adjusted with Benjamini-Hochberg method. **(B-C)** Western blot and densitometry of baseline IAP expression for **(B)** *TP53*^+/+^, *TP53^-/mut^*, and *TP53^-/-^* MOLM-13 cells or **(C)** additional *TP53*^+/+^ and *TP53^-/-^* MOLM-13 and MV-4-11 cell lines. **(D)** Upregulation of *BIRC5* mRNA in primary AML patients from the Beat AML dataset by *TP53* status (n = 404 wild-type, n = 35 mutant). Analysis via Wilcoxon test. **(E-F)** *TP53* mutant AML patients from the Munich Leukemia Laboratory dataset (n = 668 wild-type, n = 63 mutant) have significant differences in gene expression for **(E)** mRNA levels of select apoptosis genes and **(F)** *BIRC5* mRNA. Analysis via Wilcoxon test. **(G)** Patients from the MD Anderson Cancer Center cohort have upregulation of survivin measured via RPPA protein expression by *TP53* status (n = 501 wild-type, n = 153 mutant) or karyotype (n = 500 non-complex, n = 203 complex). Analysis via Wilcoxon test. **(H)** Sankey diagram showing the relationship between *BIRC5* expression levels (High vs Low) and survival outcomes (favorable vs poor) in *TP53* mutant (n = 60) and wild-type (n = 611) patients. **(I)** Boxplot showing *BIRC5* expression across different *TP53* mutation classes – missense (n = 35), frameshift (n = 9), splice site (n = 9), other (n = 7), and wild-type (n=611). Each point represents an individual patient sample. Boxes indicate interquartile range, and whiskers denote variability beyond the quartiles. Analysis via Wilcoxon rank-sum test. Flow widths represent the proportion of patients within each category. Analysis via Wilcoxon rank-sum test. Survival status was “poor” (overall survival ≤ 360 days) or “favorable” (> 360 days), with an observed cohort median of 360.5 days used as the cut-off. Thickness of flows indicates the number of patients in each survival–mutation category. For all figures, ns = no significance, *p<0.05, **p<0.01, ***p<0.001, ****p<0.0001.

### *BIRC5* is upregulated in *TP53* mutant AML patients

To determine the clinical relevance of *BIRC5* dependency in *TP53* mutant AML, we next turned to patient samples. *BIRC5* was significantly elevated in *TP53^-/mut^* tumors from the publicly available Beat AML dataset (35 *TP53^-/mut^* versus 404 *TP53^+/+^*; Figure 2D). These findings extend prior reports of elevated *BIRC5* expression in leukemia progenitor cells.^34,35^ We next validated transcriptomic signatures from newly diagnosed AML patients in the unpublished Munich Leukemia Laboratory dataset (63 *TP53^-/mut^* cases versus 668 *TP53^+/+^* cases). *TP53^-/mut^* AML showed downregulation of pro-death genes (*BAX*, *CASP2,* and *TP53*) and upregulation of pro-survival genes (*PIK3R2* and *MAPK*) (Figures 2E and S3C). Notably, *BIRC5* was among the most significantly upregulated genes in *TP53^-/mut^* tumors (Figure 2F). Reverse phase protein array analysis on primary AML samples from the MD Anderson cohort (153 *TP53^-/mut^* cases and 501 *TP53^+/+^*cases) also showed survivin upregulation at the protein level (Figure 2G). *TP53* mutant and complex karyotype AML patients had high survivin levels, which were not associated with other recurrent AML mutations, including *FLT3*, *IDH2*, *NPM1,* or *RUNX1* (Figures 2G and S3D). *TP53^-/mut^* patients had significantly worse overall survival compared to *TP53^+/+^* patients (Figure S4A). Elevated survivin levels were also associated with poorer survival, independent of *TP53* status (Figure S4B).

To further validate if the association between *BIRC5* and *TP53* mutations contributes to survival outcomes, we divided Beat AML patients into *BIRC5-*high and *BIRC5-*low groups by *TP53* status (mutant n = 60, wild-type n = 611). Almost all *BIRC5*-high patients carrying *TP53* mutations had worse survival outcomes than *TP53* wild-type patients (Figure 2H). Similarly, in patients with complex karyotypes (*TP53* mutant n = 48, *TP53* wild-type n = 147), co-presence of *TP53* mutations and high *BIRC5* led to worse survival (Figure S4C). Within the spectrum of *TP53* mutations, missense and frameshift alterations were significantly associated with increased *BIRC5* levels compared to other mutations, including splicing, insertions, and deletions (Figure 2I). Missense and frameshift mutations resulted in the most dismal survival (Figure S4D). Collectively, this highlights that *TP53* mutant patients have high levels of *BIRC5*, which contribute to poor outcomes and provide strong rationale for targeting *BIRC5* as a dependency in *TP53* mutant AML.

### Single-cell HSC/MPCs have elevated *BIRC5* in *TP53* mutant AML

Building on our findings that *BIRC5* is upregulated in *TP53* mutant AML patients, we next asked which cell subpopulations at the single-cell level contribute to high *BIRC5* expression. We performed single-cell RNA-seq on paired diagnostic and relapse samples from 22 AML patients, including 15 with *TP53* mutations and seven with *TP53* wild-type disease (Figure 3A). Using established lineage-specific gene markers, we annotated cell types to distinguish malignant leukemic blasts from non-malignant populations. After excluding non-malignant cells, we analyzed the composition of *BIRC5*-expressing malignant cells in diagnostic samples. *BIRC5* expression was enriched in mid-/late-erythroid lineage cells, plasmacytoid dendritic cells (pDC), and hematopoietic stem and multipotent progenitor cells (HSC/MPC) in *TP53* mutant cells compared to *TP53* wild-type cells. (Figures 3A and S5A).

**Figure 3.**
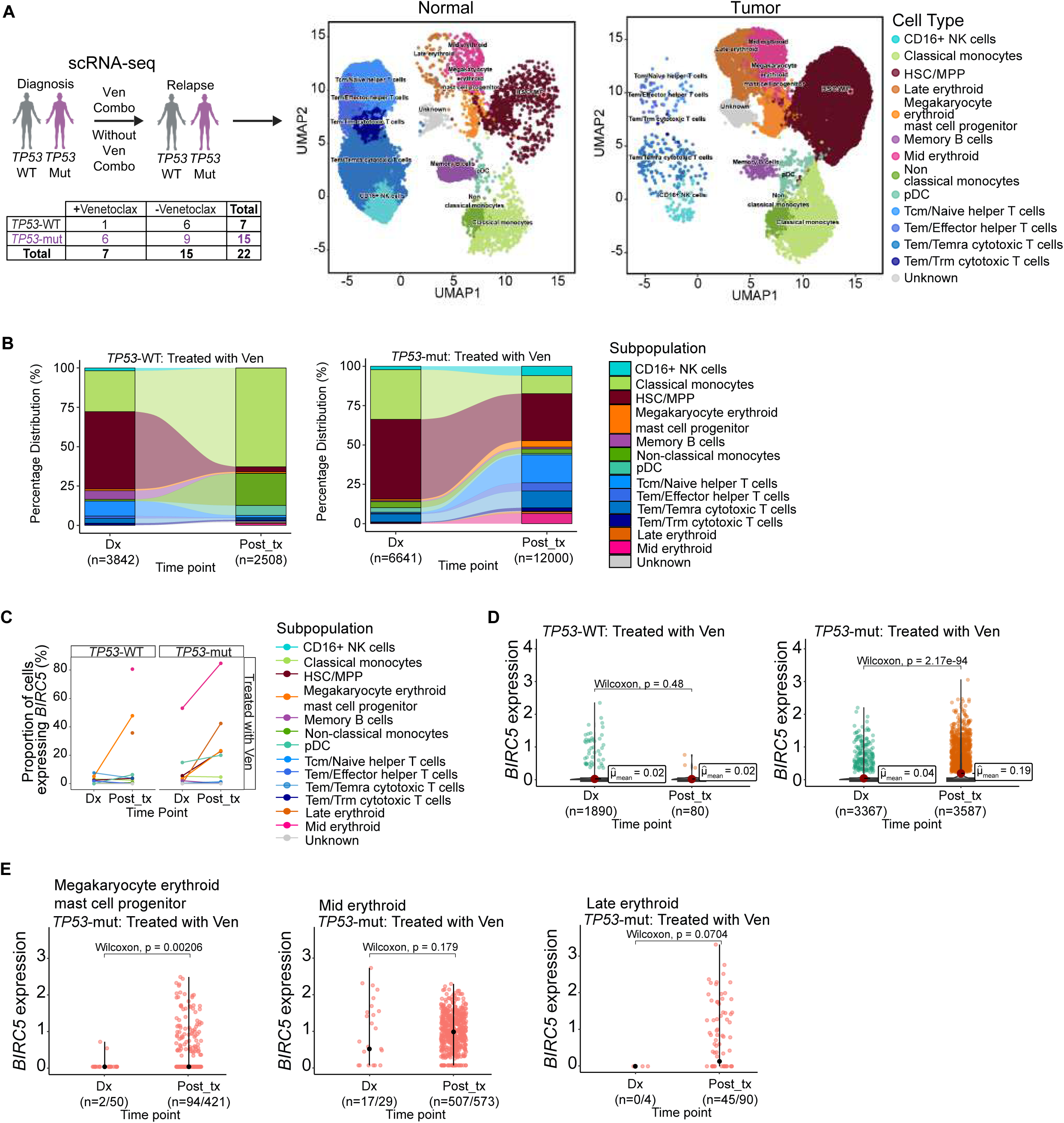
*TP53* mutant cells expressing *BIRC5^high^* clones are enriched in AML patients after relapse on venetoclax at single cell level. **(A)** Schematic and descriptive table of paired-wise samples by *TP53* mutation status and treatment type. UMAP plot visualizes the distribution of single cells based on their hematopoietic and leukemic subpopulations denoting normal vs leukemic compartments. **(B)** Sankey diagram illustrates the shift in percentage distribution of various cell subpopulations treated with venetoclax between two time points (diagnosis (Dx) and post-treatment (Post-Tx)) by *TP53* mutation status. For *TP53*-WT (n = 1 patients), single cells are n = 3842 (Dx) and n=2508 (Post-Tx). For *TP53*-Mut (n = 6 patients), single cells are n = 7748 (Dx) and n = 16899 (Post-Tx). **(C)** Proportion of cells treated with venetoclax expressing *BIRC5* (defined as *BIRC5* gene expression > 0) in various cell subpopulations between two time points (diagnosis, Dx and post-treatment, Post-Tx) by *TP53* mutation status. **(D)** Distribution of single cells based on their *BIRC5* gene expression profiles for *TP53*-WT and *TP53*-Mut cells with combination of venetoclax. For *TP53*-WT (n = 1 patients), single cells are n = 1890 (Dx) and n = 80 (Post-Tx). For *TP53*-Mut (n = 6 patients), single cells are n = 3407 (Dx) and n = 3635 (Post-Tx). **(E)** Expression levels of the *BIRC5* in HSC/MPC cell subpopulations treated with venetoclax between two timepoints Dx and Post-Tx by *TP53* mutation status for megakaryocyte erythroid mast cell, mid-erythroid, and late-erythroid populations. Wilcoxon signed rank test. ns = no significance, *p<0.05, **p<0.01, ***p<0.001, ****p<0.0001.

Longitudinal single-cell analysis of patient samples treated with and without venetoclax containing combination therapy revealed marked differences based on *TP53* status. In *TP53* wild-type patients, treatment caused significant reduction in HSC/MPC populations, indicative of favorable therapeutic response (Figures 3B and S5B). In contrast, *TP53* mutant patients retained a substantial portion of HSC/MPCs following treatment, consistent with therapy refractoriness. While *TP53* wild-type patients exhibited expansion in monocytic cell populations post-treatment, *TP53* mutant patients instead showed increased frequency of T and NK cells. Within the residual HSC/MPC compartment of *TP53* mutant patients, *BIRC5*-high clones expanded following venetoclax-based therapy and not following non-venetoclax-based regimens (Figures 3C-D and S5C-D). *BIRC5* expression significantly increased in megakaryocyte erythroid mast cells and modestly increased in mid- and late-erythroid cells after venetoclax (Figure 3E). These findings highlight *BIRC5* as a characteristic feature enriched in *TP53* mutant AML and suggest an association beween *BIRC5*-expressing HSC/MPCs and megakaryocyte erythroid cells and venetoclax resistance.

### High-throughput drug screen identifies actionable therapies that resensitize *TP53* mutant AML

Having identified *BIRC5* as a key biomarker for *TP53* mutant AML through an unbiased multiomics approach, we next sought to determine whether it truly represents a targetable vulnerability. We performed unbiased high-throughput drug screening using 289 clinically relevant agents – FDA-approved or in Phase II/III trials – in combination with VenAza across isogenic *TP53^+/+^*, *TP53^-/R248Q^*, and *TP53^-/-^* MOLM-13 cells (Figure 4A). VenAza was applied at the *TP53^+/+^* IC50 concentration and combined with each third agent at two concentrations (10 nM or 1000 nM; Figure S6A). Screen quality showed strong reproducibility across replicates and genotypes (R > 0.95; Figure S6B). As anticipated, *TP53^-/R248Q^* cells displayed reduced overall drug sensitivity compared to *TP53^+/+^* cells, with fewer compounds scoring at 10 nM (44 versus 51) and 1000 nM (79 versus 118) concentrations. Apoptosis activators, cell cycle regulators, and receptor kinase pathway modulators demonstrated the highest efficacy across genotypes (Figures 4B and S6C).

**Figure 4.**
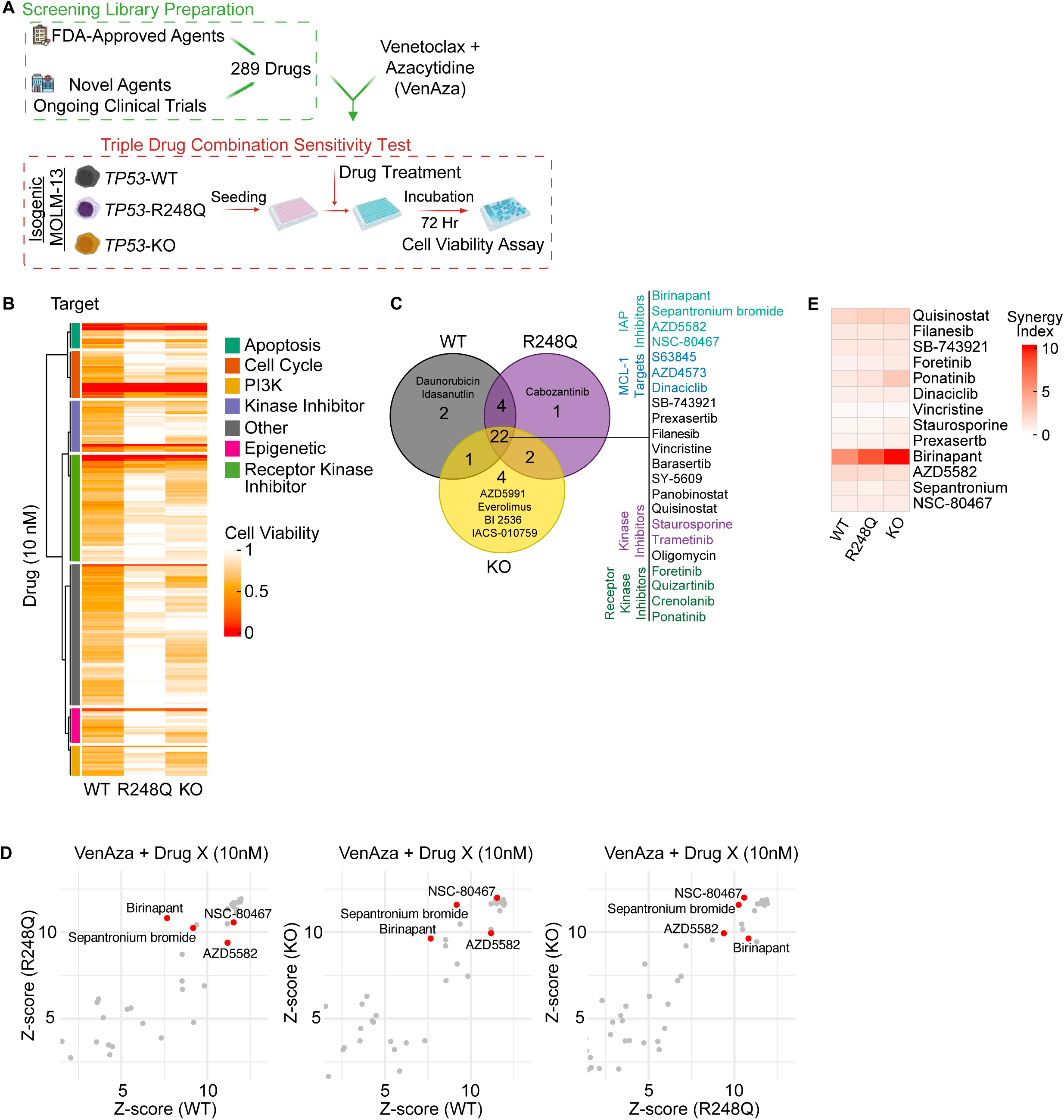
High-throughput drug screening identifies therapeutic partners for sensitizing*TP53* mutated cells to VenAza. **(A)** Schematic of high-throughput drug screening in MOLM-13 *TP53^+/+^*, *TP53^-/R248Q^*, and *TP53^-/-^* cells treated with venetoclax (3 nM), azacitidine (50 nM), with a third agent from a library of 289 clinically relevant agents. The third agent was added at concentrations of 10 nM or 1000 nM. Agents with z-score > 2 were selected. **(B)** Z-score correlation scatterplots comparing z-scores of agents at 10 nM, with comparisons between *TP53*^-/R248Q^ vs *TP53*^+/+^, and *TP53*^-/-^ vs *TP53*^+/+^. Z-score cut-off > 2 across all genotypes. **(C)** Cell viability heatmap of all agents clustered by target pathway at 10 nM concentration. **(D)** Venn diagram presenting the top 10% of scored agents from each genotype treated at 10 nM. Z-score cut-offs for *TP53^+/+^*, *TP53^-/R248Q^*, and *TP53^-/-^*cells were 4.34, 3.71, and 4.72, respectively **(E)** Synergy scores revealed that birinapant synergizes strongly with VenAza in MOLM-13 *TP53^+/+^*, *TP53^-/R248Q^*, and *TP53^-/-^*cells. For all figures, the experiments were repeated twice. One biological replicate is shown with three technical replicates.

Among the top 10% of scoring compounds (29 hits), the majority (22/29) were effective across all genotypes, suggesting *TP53*-independent anti-leukemic activity (Figure 4C and S6D). Top scoring agents were histone deacetylases (HDAC) inhibitors (quisinostat, panobinostat), cell cycle regulators (dinaciclib, prexasertib, filanesib), and apoptosis modulators (MCL-1 inhibitors, S63845 and AZD4573). As expected, daunorubicin and idasanutlin scored selectively in *TP53^+/+^*cells.^36^ Survivin inhibitors (sepantronium bromide, NSC-80467) and IAP antagonists (birinapant, AZD5582) scored preferentially in *TP53^-/R248Q^*and *TP53^-/-^* cells, in line with our CRISPR screen (Figures 4D and S6E). Further validation using seven dose-response concentrations revealed thirteen agents with potent monotherapy activity (IC50 < 20 nM), including IAP inhibitors (Figure S6F). Among these, birinapant exhibited the strongest synergy when combined with VenAza (Figure 4E). By combining complementary CRISPR and high-throughput drug screenings, we not only validate *BIRC5* as a dependency but also identify IAP proteins more broadly as actionable vulnerabilities to overcome therapy resistance in *TP53* mutant AML.

### Survivin and IAP inhibitors/degrader restore sensitivity to VenAza in *TP53* mutant AML

We evaluated the therapeutic potential of top scoring survivin (sepantronium bromide, NSC-80467) and IAP (birinapant, AZD5582) inhibitors in *TP53* mutant AML. As single agents, AZD5582, birinapant, sepantronium bromide, and NSC-80467 demonstrated potent cytotoxicity against both *TP53^-/R248Q^* and *TP53^-/-^* cells (Figure S7A). The cytotoxic effects of birinapant, sepantronium bromide, and NSC-80467 extended to *TP53^-/R175H^*, *TP53^-/R273H^*, *TP53^-/R282W^*, *TP53^- /Y220C^*, and *TP53^-/M237I^*cells. This supports broad applicability of these agents despite heterogeneous survivin expression (seen in Figure 2B) and genetically diverse *TP53* mutant backgrounds (Figures S7B-C). Combining VenAza with birinapant (VenAza-Bir) resulted in over a 10-fold reduction in venetoclax IC50 in *TP53^-/R248Q^* cells while overall inducing apoptosis across the three genotypes (Figures 5A-B).

**Figure 5.**
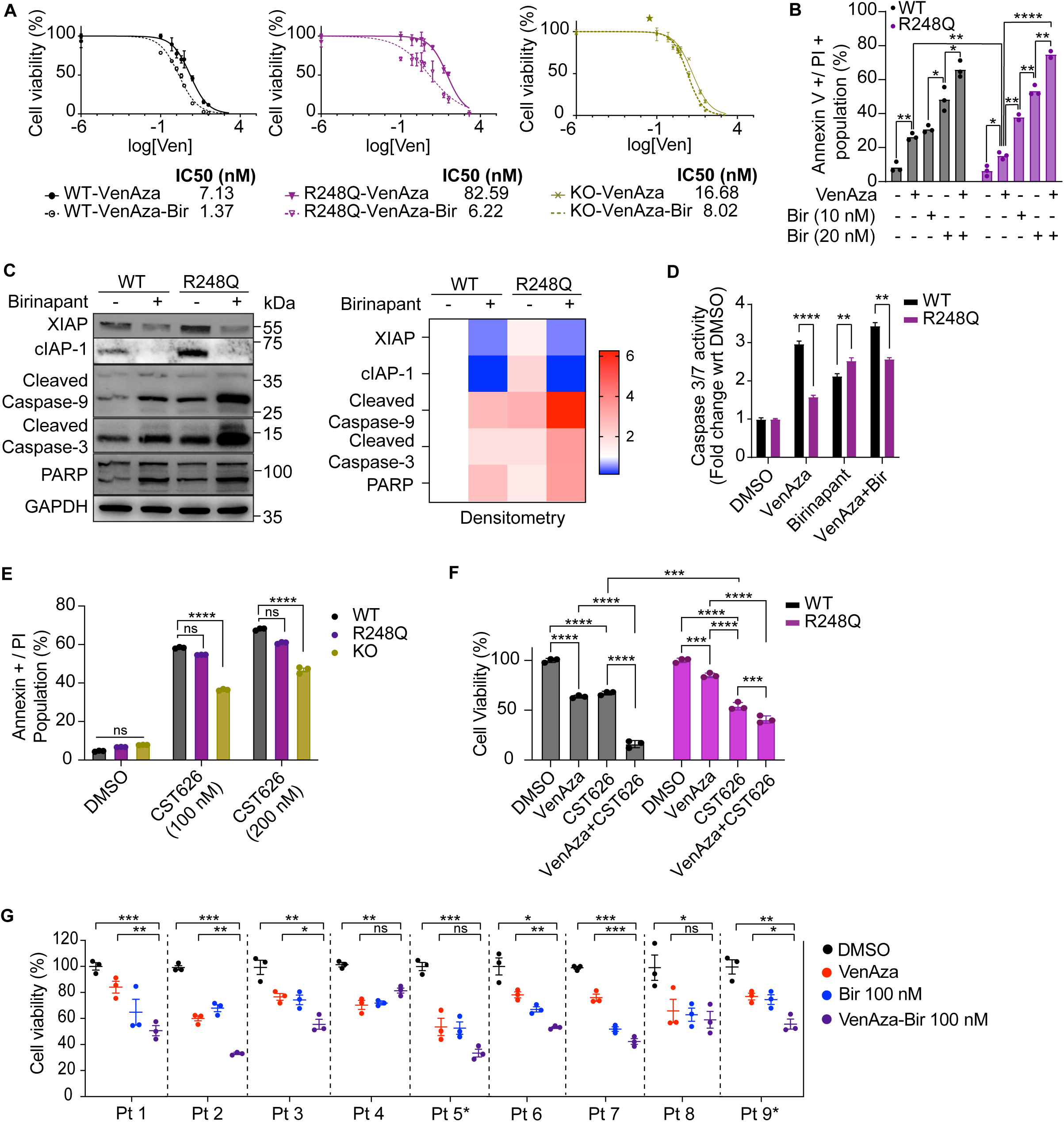
Survivin or IAP inhibitors resensitize *TP53* mutated AML cells to VenAza. **(A)** Cell viability in isogenic MOLM-13 *TP53^+/+^*, *TP53^-/R248Q^*, and *TP53^-/-^* cells treated with birinapant (20 nM) in combination with VenAza for 72 hours. **(B)** Annexin V-APC and PI assay shows increased cell death populations in *TP53*^+/+^ and *TP53*^-/R248Q^ cells treated with VenAza, birinapant, and VenAza plus birinapant (10 nM, 20 nM) for 72 hours. **(C)** Western blot with densitometry of cleaved caspase-9, cleaved caspase-3 and total caspase-3 after treatment of MOLM-13 *TP53*^+/+^ and *TP53*^-/R248Q^ with VenAza across different timepoints. **(D)** Caspase-3/7 activity changes in isogenic MOLM-13 cells treated with DMSO, VenAza, birinapant (20 nM), or VenAza-Bir for 24 hours. **(E)** Annexin+ PI levels for MOLM-13 *TP53^+/+^*, *TP53^-/R248Q^*, and *TP53^-/-^* cells treated with IAP degrader, CST626, at 100 nM and 200 nM shows increased apoptosis. **(F)** Combination of VenAza plus CST626 (100 nM) increases cytotoxicity in *TP53^+/+^*and *TP53^-/R248Q^* cells compared to VenAza or CST626 alone. **(G)** Cell viability in nine primary *TP53* mutant AML patient cells treated DMSO, VenAza, birinapant (100 nM), or VenAza-Bir for 72 hours shows increased sensitivity to triplet therapy. For patients with an *, birinapant was 500 nM. For all figures, treatment with VenAza: venetoclax 3 nM, azacitidine 50 nM. ns = no significance, *p<0.05, **p<0.01, ***p<0.001, ****p<0.0001.

Since *BIRC5*-KO increased caspase activation, we next investigated whether IAP inhibition could restore apoptosis by re-engaging the caspase cascade. Consistent with its function as a SMAC mimetic, birinapant reduced XIAP and cIAP-1 protein levels in *TP53^+/+^* and *TP53^-/R248Q^* cells (Figure 5C). In *TP53^-/R248Q^* cells, birinapant induced robust cleavage of caspase-9, caspase-3/7, and PARP, along with a rise in caspase-3/7 enzymatic activity – effects not observed with VenAza alone (Figures 5C-D). Birinapant also triggered mitochondrial membrane potential across all genotypes, indicating restored intrinsic apoptotic and caspase signaling (Figure S7D). Treatment with IAP degrader CST626 (targets XIAP, cIAP-1 and cIAP-2^37^) increased apoptosis across all three genotypes (Figure 5E). Combination of VenAza plus CST626 also showed impressive cytotoxic effects compared to VenAza or CST626 alone (Figure 5F). Next, we treated nine *TP53* mutant AML primary tumors with VenAza, birinapant, and VenAza plus birinapant (VenAza-Bir). VenAza-Bir caused increased cytotoxicity compared to either VenAza or birinapant alone (statistically significant for 6/9 patients), suggesting translational potential for IAP inhibitors (Figure 5G).

Survivin serves dual roles – promotes mitotic progression and inhibits apoptosis by blocking caspase activation.^38^ We next asked by which mechanism survivin inhibitors restored VenAza sensitivity. Both survivin inhibitors rescued caspase-3/7 activity in *TP53^+/+^*, *TP53^-/R248Q^*, and *TP53^-/-^*cells (Figure S8A-B). Sepantronium bromide also increased annexin V/PI staining and induced accumulation of cells in the sub-G1 phase, consistent with elevated levels of apoptosis (Figures S8C-D). However, treatment with sepantronium bromide did not result in post-mitotic arrest or spindle defects, suggesting that its effects are not due to disruption of mitotic entry (Figures S8E). Together, these findings demonstrate that *BIRC5* dependency in *TP53* mutant AML is mechanistically driven by its anti-apoptotic function – specifically suppression of post-mitochondrial caspase activity – and that targeting survivin or related IAP family members can restore caspase-dependent apoptosis to overcome VenAza resistance.

### Survivin and IAP inhibitors cause leukemia inhibition *in vivo* in TP53 mutant AML models

We next validated the *in vivo* efficacy of survivin and IAP inhibitors in combination with VenAza (Figure 6A). To balance efficacy with tolerability, we titrated venetoclax to a lower concentration (30 mg/kg/day) previously shown to be efficacious.^39–42^ As expected, birinapant monotherapy was ineffective and showed no survival benefit in *TP53^-/R248Q^* mice compared to control-treated *TP53^+/+^* mice (median survival 20 versus 19 days) (Figure S9A). Thus, we simulated a realistic clinical scenario comparing VenAza-Bir to VenAza in *TP53^+/+^* and *TP53^- /R248Q^* mice. We first conducted a proof of concept experiment to test if triplet therapy in *TP53^- /R248Q^* mice exceeded therapeutic benefits of VenAza in *TP53^+/+^* mice. After five days post-engraftment, VenAza-Bir significantly extended survival for *TP53^-/R248Q^* mice (median 44.5 days) compared to VenAza alone in *TP53^+/+^* mice (median 26 days) (Figure S9B).

**Figure 6.**
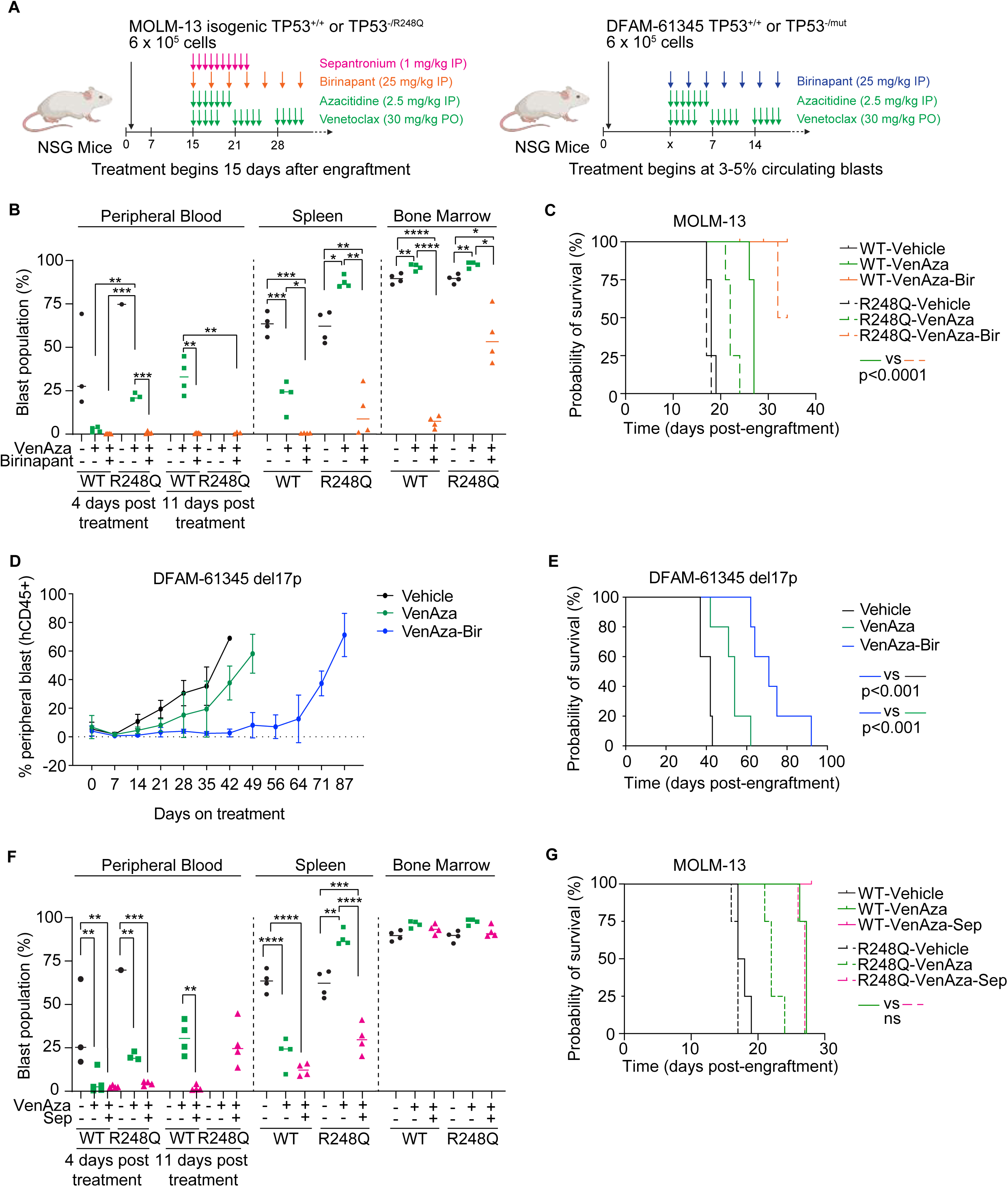
IAP or survivin inhibitors cause *in vivo* leukemia inhibition in TP53 mutant AML models. **(A)** Schematic of *in vivo* study where NSG mice (n = 30) were engrafted with either isogenic MOLM-13 *TP53*^+/+^ or *TP53*^-/R248Q^ cells. At 15 days post-transplant, mice were further split into six groups and were treated with either vehicle, VenAza, or VenAza plus birinapant (VenAza-Bir). **(B)** Blast count percentages are reduced in the peripheral blood of *TP53*^+/+^ or *TP53*^-/R248Q^ Molm-13 isogenic mice 4 days and 11 days post-treatment with VenAza-Bir. Blast count percentages in spleen and bone marrow are reduced at the end point. **(C)** Survival percentage of *TP53*^+/+^ or *TP53*^-/R248Q^ Molm-13 isogenic mice by treatment group post-transplant shows increased survival in mice receiving VenAza-Bir. **(D)** *TP53* null (del 17p) DFAM-61345 PDX from chemotherapy relapsed AML patient were transplanted in NSG mice. At 3-5% circulating blasts, mice were further split into 3 groups with either vehicle, VenAza, or VenAza-Bir. Blast count percentages are substantially reduced in the peripheral blood with VenAza-Bir. **(E)** Survival percentage of DFAM-61345 PDX mice by treatment group post-transplant shows increased survival in mice receiving VenAza-Bir relative to VenAza or vehicle. **(F)** Blast count percentages are reduced in the peripheral blood of *TP53*^+/+^ or *TP53*^-/R248Q^ Molm-13 isogenic mice 4 days and 11 days post-treatment with VenAza-Sep. Blast count percentages in spleen and bone marrow are reduced at the end point. **(G)** Survival percentage of *TP53*^+/+^ or *TP53*^-/R248Q^ Molm-13 isogenic mice by treatment group post-transplant shows increased survival in mice receiving VenAza-Sep. For all figures: venetoclax 30 mg/kg PO, azacitidine 2.5 mg/kg IP, birinapant 25 mg/kg IP, sepantronium bromide 1 mg/kg IP.

Having established superior efficacy in this early-intervention model, we next assessed VenAza-Bir in a highly-aggressive setting. Therapy was initiated at 15 days post-engraftment, when mice displayed partial hind-limb paralysis and control mice had succumbed to disease (median 18 days). VenAza-Bir treatment markedly reduced leukemic burden in circulation, spleen, and bone marrow (Figure 6B). This enhanced disease control translated to improved survival for *TP53^-/R248Q^*mice treated with VenAza-Bir (median 33 days) compared to VenAza (median 22 days) (Figure 6C). Triplet therapy in *TP53^-/R248Q^* mice also exceeded standard of care VenAza in *TP53^+/+^*mice (median 27 days). In the AML patient-derived xenograft model DFAM-61345 harboring *TP53* deletion (del 17p) established from a chemotherapy-resistant patient, VenAza-Bir treatment significantly suppressed circulating leukemic blast counts and extended median survival to 72 days compared to 54 days in mice treated with VenAza alone (Figures 6D-E). Importantly, treatments were well-tolerated, with no significant drops in body weight and no evident toxicity in lung, liver, heart, or kidney tissues (Figures S9C-D).

VenAza plus sepantronium bromide (VenAza-Sep) also showed anti-leukemic efficacy in *TP53^-/R248Q^*mice, significantly reducing leukemic burden in peripheral blood and spleen but not bone marrow (Figure 6F). Reduced activity in bone marrow may be attributable to the short plasma half-life of sepantronium bromide (approximately 1-2 hours)^43^, potentially causing suboptimal drug penetration. Nevertheless, VenAza-Sep treatment modestly extended survival in *TP53^-/R248Q^* mice (median 27 days) compared to VenAza (median 22 days) and matched the survival of *TP53^+/+^* mice treated with VenAza (median 27 days) (Figure 6G). Collectively, this supports IAP inhibition – particularly with birinapant and to a lesser extent sepantronium bromide – as robust and clinically promising triplet strategy to replace existing standard of care VenAza in *TP53* mutant AML.

### Survivin inhibitors overcome chemotherapy resistance in *TP53* deficient solid cancers

To test the broader implications of *BIRC5* targeting across *TP53* mutant malignancies, we next turned to solid cancers. In the pan-cancer TCGA cohort, *BIRC5* was upregulated in 17 out of 25 different *TP53* mutant cancer types (Figure 7A). We then asked whether the mechanism of therapy resistance observed in AML^22^ extended to other *TP53* deficient tumors. We evaluated responses to cytotoxic chemotherapy in two *TP53^-/-^* solid cancers: colorectal cancer (CRC; HCT-116) treated with 5-flurouracil (5-FU) and triple-negative breast cancer (TNBC; CAL-51) treated with doxorubicin (Dox). *TP53^-/-^*CRC and TNBC cells were more chemoresistant (Figures 7B and S10A). *TP53* loss consistently impaired caspase-3/7 activation in response to chemotherapy in CRC and TNBC cells too, mirroring resistance phenotypes in AML (Figure 7C). In *TP53^-/-^*HCT-116 cells, survivin and XIAP were upregulated, implicating IAPs as a shared dependency across both hematological and solid tumors lacking functional p53 (Figure 7D).

**Figure 7.**
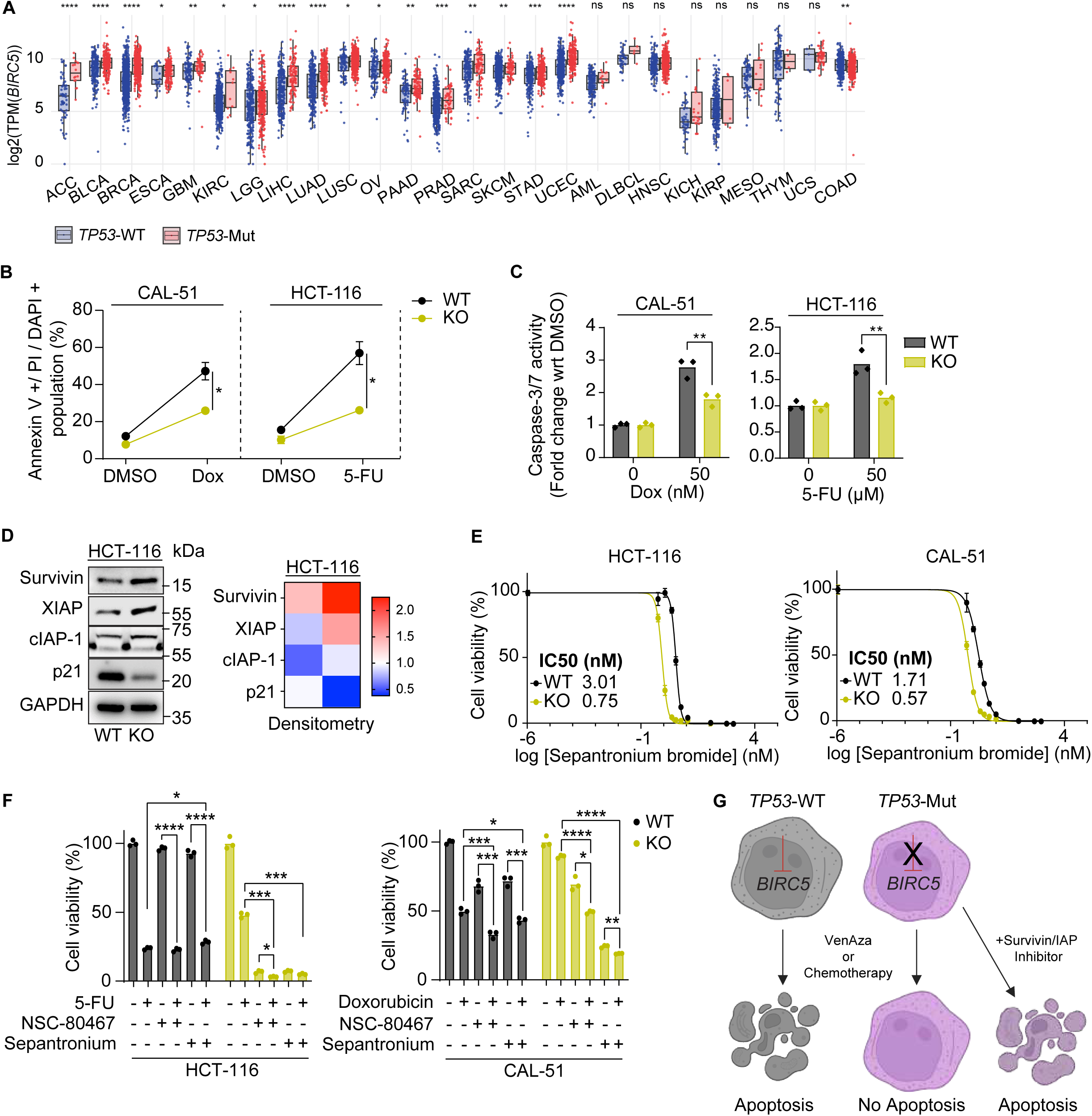
*TP53* mutant solid cancers exhibit *BIRC5*/survivin dependency. **(A)** Upregulation of *BIRC5* mRNA obtained from RNA-seq of individual patients from 17 of 25 cancers in the TCGA database. Analysis via Wilcoxon test. **(B)** Annexin V-APC and PI assay for *TP53^-/-^*triple negative breast cancer (CAL-51) and colorectal (HCT-116) cell lines treated with either standard of care doxorubicin (10 nM, 100 nM) or 5-fluorouracil (5-FU; 50 mM, 100 mM), respectively, show increased cell death populations. **(C)** Caspase-3/7 activity is diminished in *TP53^-/-^* CAL-51 and HCT-116 cells treated with doxorubicin or 5-FU. **(D)** Western blot and densitometry of HCT-116 cells show increased survivin and XIAP expression *TP53^-/-^*cells at baseline. **(E)** Cell viability plots for *TP53*^+/+^ and *TP53*^-/-^ CAL-51 and HCT-116 cells treated for 72 hours with sepantronium bromide alone. **(F)** Cell viability for CAL-51 and HCT-116 cells treated with doxorubicin or 5-FU plus a sepantronium bromide or NSC-80467. **(G)** Schematic of survivin/IAP inhibitors restoring apoptosis in *TP53* mutant cancers. For all figures unless otherwise specified, treatment with doxorubicin 50 nM, 5-FU 50 mM, sepantronium bromide 3 nM, and NSC-80467 10 nM. ns = no significance, *p<0.05, **p<0.01, ***p<0.001, ****p<0.0001.

We next exploited survivin/IAP upregulation to target *TP53^-/-^*CRC and TNBC cells. Survivin inhibitors had greater or comparable monotherapy efficacy in *TP53^-/-^* HCT-116 and CAL-51 cells compared to *TP53^+/+^* cells (Figures 7E and S10B). Strikingly, co-treatment with survivin inhibitors and 5-FU or doxorubicin restored chemosensitivity in *TP53^-/-^* HCT-116 and CAL-51 cells (Figure 7F). In contrast, birinapant failed to augment chemotherapy efficacy in *TP53^-/-^* HCT-116 and CAL-51 cells (Figure S10C). These findings highlight a key distinction between AML and solid tumors – where IAP inhibition sensitized *TP53* mutant AML to VenAza, survivin inhibition appears to be a consistently effective modality for hematological and solid cancers with *TP53* mutations and deletions (Figure 7G).

## DISCUSSION

Clinically, most *TP53* mutant patients relapse within six months of initiating frontline VenAza therapy^7^, highlighting the urgent need for more effective strategies. Ominously, recent clinical trials in *TP53* mutant AML with eprenetapopt (p53 activator) plus azacitidine and magrolimab (anti-CD47 antibody) plus azacitidine have been met with disappointment.^44–47^ This underscores the limitations of strategies that rely on restoring or bypassing upstream p53 function without addressing downstream apoptotic blockades. In contrast, our study demonstrates that survivin and IAPs are critical, druggable mediators of post-mitochondrial resistance. Combination of survivin and IAP inhibitor with VenAza significantly enhanced leukemia suppression *in vivo*, outperforming VenAza alone in both *TP53* mutant and wild-type settings. This approach has major translational implications for highly-resistant *TP53* mutant AML.

Prior studies have focused on restoring canonical p53 signaling through transcriptional activation of wild-type target genes using p53 activators, BAX activators, and STING agonists to overcome therapy resistance.^25,48,49^ Here, we undertook an unbiased approach to determine whether one downstream target stands out from the rest as a core driver of therapy resistance. We unveiled *BIRC5* as one such functional dependency in *TP53* mutant AML. Importantly, our comprehensive multiomics (CRISPR and high-throughput drug screens) address key limitations inherent to either method alone: (1) gene dependencies identified by CRISPR may not always be pharmacologically actionable, (2) drugs that act through multi-target mechanisms may be missed by single-gene knockout screens, and (3) narrow strategies may overlook other clinically relevant drugs.

Although *BIRC5* overexpression in cancer has been recognized^50^, our findings highlight the importance of genomic context in regulating *BIRC5* expression. Among recurrent AML-associated mutations (*FLT3, IDH2, NPM1, RUNX1*), only *TP53* mutations and complex karyotypes showed consistent *BIRC5* upregulation. While heterogeneity in *BIRC5* expression levels was observed in different *TP53* mutant and non-mutant cases, elevated *BIRC5* was consistently associated with unfavorable outcomes. Functionally, *BIRC5* deletion in AML also restored caspase-9 and -3/7 activity and downregulated other IAPs, implicating survivin as the central post-MOMP regulator for blocking apoptosis. However, survivin’s direct role in caspase inhibition is controversial due to lack of a C-terminal ring finger domain.^28,31^ Some studies have proposed indirect modes of action – stabilization of XIAP to support caspase-9 binding, formation of a survivin-HBXIP complex that binds procaspase-9, or SMAC antagonization.^28^ While most studies have focused on *TP53* deletions, there is also growing evidence that *TP53* mutations confer distinct biological phenotypes and clinical outcomes^51^, which we capture here. Across six recurrent AML-associated *TP53* missense mutations, caspase blockade was a conserved phenomenon to evade therapy-induced apoptosis.

*BIRC5* is a targetable vulnerability to overcome therapy resistance in *TP53* mutant AML, and its dual role in mitosis and apoptosis make it a uniquely promising target. In *TP53* mutant AML, survivin primarily functions as an IAP to block caspase activation and prevent cell death, which can be overcome by pharmacologic inhibition. Despite failure of survivin inhibition to cause mitotic arrest in *TP53* mutant AML, cells were vulnerable to other G2/M transition modulators identified from our drug screen, including aurora-B kinase inhibitor, barasertib, and kinesin spindle protein inhibitor, filanesib. This sensitivity may arise because *TP53* mutant cells, while able to bypass G1/S checkpoints, become increasingly dependent on successful mitotic progression and upregulated survivin to support this process. This broad activity underscores therapeutic versatility of survivin inhibitors as combination partners.

Personalizing therapy by stratifying patients to the most effective treatments is one of the most important steps for improving clinical outcomes in *TP53* mutant AML. Our longitudinal single-cell RNA-seq analysis revealed selective enrichment of *BIRC5* expression within therapy-resistant HSC/MPCs following venetoclax-based treatment. *BIRC5* was also upregulated in erythroid populations, which supports clinical observations that erythroid leukemia patients have a higher burden of *TP53* mutations. This finding implies that in *TP53* mutant patients, *BIRC5^high^* clones expand during treatment and guides the rational application of survivin and IAP inhibitor therapies in highly-aggressive *TP53* mutant and deficient cancers.

## Acknowledgments

S.B. acknowledges support from the Singapore Ministry of Education (MOE) Academic Research Fund (AcRF) Tier 2 (MOE-000573-00), NUS-Paris University Joint Grant, and Leukemia Research Foundation New Investigator Research Program. S.B. is a recipient of the EMBO Global Investigator Award. A.M.M. acknowledges support from the Singapore International Graduate Award (SINGA). A.P. acknowledges support by ERC CoG (DynAML, 101088563) and LNCC (EL2024-PUISSANT) grants. J.K. acknowledges support from the José Carreras Leukämie-Stiftung (project DJCLS 07 R/2024), the Deutsche Forschungsgesellschaft (DFG KR 3886/8-1), and the Deutsche Konsortium für Translationale Krebsforschung (DKTK).

## Author Contributions

S.B. conceptualized the study, designed the experiments, and wrote the manuscript. A.M.M. and F.Q.L. conceptualized and designed experiments, performed experiments, analyzed data, and contributed to manuscript construction. Y.M. performed high-throughput drug screening and validation in isogenic AML cell lines. E.A.O. designed figures and wrote the manuscript. C.G.T.C., Y.W., J.Y.M.T., N.C., and K.S.B. performed experiments and analyzed data. S.J. performed BH3 profiling and DBP experiments and analyzed data. X.X.L. and A.N.M. analyzed and plotted data. V.S. performed proteomics experiments, analyzed data, and critically reviewed the manuscript. C.W. generated *TP53* knockout AML cell lines. P.M. performed mass spectrometry proteomics and analyzed data. S.G., K.T., and S.D. analyzed transcriptomics, *BIRC5* expression, and survival from the Beat AML dataset. B.D.B. and S.M.K. designed survival experiments, measured survivin levels in the MD Anderson patient dataset, and analyzed data. T.H. provided bulk transcriptomic analysis of the MLL patient dataset. A.P. supervised and analyzed with C.L. bulk RNAseq in isogenic AML cell lines. P.G.M. critically reviewed the manuscript. J.K. designed and supervised experiments, analyzed proteomics data, and critically reviewed the manuscript. E.A. and H.A. analyzed single cell and bulk data from the MLL dataset and MD Anderson Moon Shots dataset. K.I. provided expertise regarding acquisition of *BIRC5* upregulation and critically reviewed the manuscript. M.A. supervised experiments on the MD Anderson dataset, provided patient samples, and contributed to single cell/bulk RNAseq and proteomics experiments. J.T. critically reviewed the manuscript.

## REFERENCES

1. Toma-Jonik A, Vydra N, Janus P, Widłak W. Interplay between HSF1 and p53 signaling pathways in cancer initiation and progression: non-oncogene and oncogene addiction. Cell Oncol Dordr. 2019;42(5):579–589. doi:10.1007/s13402-019-00452-0

2. Kastenhuber ER, Lowe SW. Putting p53 in Context. Cell. 2017;170(6):1062–1078. doi:10.1016/j.cell.2017.08.028

3. Barbosa K, Li S, Adams PD, Deshpande AJ. The role of TP53 in acute myeloid leukemia: Challenges and opportunities. Genes Chromosomes Cancer. 2019;58(12):875–888. doi:10.1002/gcc.22796

4. Papaemmanuil E, Gerstung M, Bullinger L, et al. Genomic Classification and Prognosis in Acute Myeloid Leukemia. N Engl J Med. 2016;374(23):2209–2221. doi:10.1056/NEJMoa1516192

5. Lindsley RC, Saber W, Mar BG, et al. Prognostic Mutations in Myelodysplastic Syndrome after Stem-Cell Transplantation. N Engl J Med. 2017;376(6):536–547. doi:10.1056/NEJMoa1611604

6. Feurstein S, Rücker FG, Bullinger L, et al. Haploinsufficiency of ETV6 and CDKN1B in patients with acute myeloid leukemia and complex karyotype. BMC Genomics. 2014;15(1):784. doi:10.1186/1471-2164-15-784

7. Pollyea DA, Pratz KW, Wei AH, et al. Outcomes in Patients with Poor-Risk Cytogenetics with or without TP53 Mutations Treated with Venetoclax and Azacitidine. Clin Cancer Res Off J Am Assoc Cancer Res. 2022;28(24):5272–5279. doi:10.1158/1078-0432.CCR-22-1183

8. DiNardo CD, Pratz K, Pullarkat V, et al. Venetoclax combined with decitabine or azacitidine in treatment-naive, elderly patients with acute myeloid leukemia. Blood. 2019;133(1):7–17. doi:10.1182/blood-2018-08-868752

9. Konopleva M, Letai A. BCL-2 inhibition in AML: an unexpected bonus? Blood. 2018;132(10):1007–1012. doi:10.1182/blood-2018-03-828269

10. Wei AH, Strickland SA, Hou JZ, et al. Venetoclax Combined With Low-Dose Cytarabine for Previously Untreated Patients With Acute Myeloid Leukemia: Results From a Phase Ib/II Study. J Clin Oncol Off J Am Soc Clin Oncol. 2019;37(15):1277–1284. doi:10.1200/JCO.18.01600

11. DiNardo CD, Jonas BA, Pullarkat V, et al. Azacitidine and Venetoclax in Previously Untreated Acute Myeloid Leukemia. N Engl J Med. 2020;383(7):617–629. doi:10.1056/NEJMoa2012971

12. Kim K, Maiti A, Loghavi S, et al. Outcomes of TP53-mutant acute myeloid leukemia with decitabine and venetoclax. Cancer. 2021;127(20):3772–3781. doi:10.1002/cncr.33689

13. Daver N, Wei AH, Pollyea DA, Fathi AT, Vyas P, DiNardo CD. New directions for emerging therapies in acute myeloid leukemia: the next chapter. Blood Cancer J. 2020;10(10):107. doi:10.1038/s41408-020-00376-1

14. Yu J, Wang Z, Kinzler KW, Vogelstein B, Zhang L. PUMA mediates the apoptotic response to p53 in colorectal cancer cells. Proc Natl Acad Sci U S A. 2003;100(4):1931–1936. doi:10.1073/pnas.2627984100

15. Oda E, Ohki R, Murasawa H, et al. Noxa, a BH3-only member of the Bcl-2 family and candidate mediator of p53-induced apoptosis. Science. 2000;288(5468):1053–1058. doi:10.1126/science.288.5468.1053

16. Miyashita T, Harigai M, Hanada M, Reed JC. Identification of a p53-dependent negative response element in the bcl-2 gene. Cancer Res. 1994;54(12):3131–3135.

17. Wu Y, Mehew JW, Heckman CA, Arcinas M, Boxer LM. Negative regulation of bcl-2 expression by p53 in hematopoietic cells. Oncogene. 2001;20(2):240–251. doi:10.1038/sj.onc.1204067

18. Tait SWG, Parsons MJ, Llambi F, et al. Resistance to caspase-independent cell death requires persistence of intact mitochondria. Dev Cell. 2010;18(5):802–813. doi:10.1016/j.devcel.2010.03.014

19. Chipuk JE, Kuwana T, Bouchier-Hayes L, et al. Direct activation of Bax by p53 mediates mitochondrial membrane permeabilization and apoptosis. Science. 2004;303(5660):1010–1014. doi:10.1126/science.1092734

20. Chipuk JE, Bouchier-Hayes L, Kuwana T, Newmeyer DD, Green DR. PUMA couples the nuclear and cytoplasmic proapoptotic function of p53. Science. 2005;309(5741):1732–1735. doi:10.1126/science.1114297

21. Leu JIJ, Dumont P, Hafey M, Murphy ME, George DL. Mitochondrial p53 activates Bak and causes disruption of a Bak-Mcl1 complex. Nat Cell Biol. 2004;6(5):443–450. doi:10.1038/ncb1123

22. Mamdouh AM, Olesinski EA, Lim FQ, et al. TP53 mutations drive therapy resistance via post-mitochondrial caspase blockade. bioRxiv. Preprint posted online August 31, 2025:2025.08.28.672283. doi:10.1101/2025.08.28.672283

23. Aubrey BJ, Kelly GL, Janic A, Herold MJ, Strasser A. How does p53 induce apoptosis and how does this relate to p53-mediated tumour suppression? Cell Death Differ. 2018;25(1):104–113. doi:10.1038/cdd.2017.169

24. Boettcher S, Miller PG, Sharma R, et al. A dominant-negative effect drives selection of TP53 missense mutations in myeloid malignancies. Science. 2019;365(6453):599–604. doi:10.1126/science.aax3649

25. Diepstraten ST, Yuan Y, La Marca JE, et al. Putting the STING back into BH3-mimetic drugs for TP53-mutant blood cancers. Cancer Cell. 2024;42(5):850–868.e9. doi:10.1016/j.ccell.2024.04.004

26. Doench JG, Fusi N, Sullender M, et al. Optimized sgRNA design to maximize activity and minimize off-target effects of CRISPR-Cas9. Nat Biotechnol. 2016;34(2):184–191. doi:10.1038/nbt.3437

27. Fulda S, Vucic D. Targeting IAP proteins for therapeutic intervention in cancer. Nat Rev Drug Discov. 2012;11(2):109–124. doi:10.1038/nrd3627

28. Marusawa H, Matsuzawa S, Welsh K, et al. HBXIP functions as a cofactor of survivin in apoptosis suppression. EMBO J. 2003;22(11):2729–2740. doi:10.1093/emboj/cdg263

29. Dohi T, Okada K, Xia F, et al. An IAP-IAP Complex Inhibits Apoptosis *. J Biol Chem. 2004;279(33):34087–34090. doi:10.1074/jbc.C400236200

30. Hoffman WH, Biade S, Zilfou JT, Chen J, Murphy M. Transcriptional repression of the anti-apoptotic survivin gene by wild type p53. J Biol Chem. 2002;277(5):3247–3257. doi:10.1074/jbc.M106643200

31. Bratton SB, Walker G, Srinivasula SM, et al. Recruitment, activation and retention of caspases-9 and -3 by Apaf-1 apoptosome and associated XIAP complexes. EMBO J. 2001;20(5):998. doi:10.1093/emboj/20.5.998

32. Murphy M, Hinman A, Levine AJ. Wild-type p53 negatively regulates the expression of a microtubule-associated protein. Genes Dev. 1996;10(23):2971–2980. doi:10.1101/gad.10.23.2971

33. Hao Y, Sekine K, Kawabata A, et al. Apollon ubiquitinates SMAC and caspase-9, and has an essential cytoprotection function. Nat Cell Biol. 2004;6(9):849–860. doi:10.1038/ncb1159

34. Carter BZ, Qiu Y, Huang X, et al. Survivin is highly expressed in CD34+38− leukemic stem/progenitor cells and predicts poor clinical outcomes in AML. Blood. 2012;120(1):173–180. doi:10.1182/blood-2012-02-409888

35. Carter BZ, Wang RY, Schober WD, Milella M, Chism D, Andreeff M. Targeting Survivin expression induces cell proliferation defect and subsequent cell death involving mitochondrial pathway in myeloid leukemic cells. Cell Cycle Georget Tex. 2003;2(5):488–493.

36. Ventura A, Kirsch DG, McLaughlin ME, et al. Restoration of p53 function leads to tumour regression in vivo. Nature. 2007;445(7128):661–665. doi:10.1038/nature05541

37. Steinebach C, Bricelj A, Murgai A, et al. Leveraging Ligand Affinity and Properties: Discovery of Novel Benzamide-Type Cereblon Binders for the Design of PROTACs. J Med Chem. 2023;66(21):14513–14543. doi:10.1021/acs.jmedchem.3c00851

38. Li F, Ambrosini G, Chu EY, et al. Control of apoptosis and mitotic spindle checkpoint by survivin. Nature. 1998;396(6711):580–584. doi:10.1038/25141

39. Zha J, Zhou H, Zhong M, et al. Preclinical Studies and Phase II Trial of Venetoclax in Combination with Chidamide and Azacitidine in Relapsed/Refractory Acute Myeloid Leukemia. Blood. 2022;140(Supplement 1):3292–3293. doi:10.1182/blood-2022-165261

40. M A, A S, T S, T M. Shorter duration of venetoclax administration to 14 days has same efficacy and better safety profile in treatment of acute myeloid leukemia. Ann Hematol. 2023;102(3). doi:10.1007/s00277-023-05102-y

41. Borate U, Huang Y, Zeidner JF, et al. A Randomized Phase 2 Trial of 28-Day (Arm A) Versus 14-Day (Arm B) Schedule of Venetoclax + Azacitidine in Newly Diagnosed Acute Myeloid Leukemia Patients ≥ 60 Years. Blood. 2024;144(Supplement 1):2907.2. doi:10.1182/blood-2024-210967

42. Garciaz S, Saillard C, Hicheri Y, Hospital MA, Vey N. Venetoclax in Acute Myeloid Leukemia: Molecular Basis, Evidences for Preclinical and Clinical Efficacy and Strategies to Target Resistance. Cancers. 2021;13(22):5608. doi:10.3390/cancers13225608

43. Gholizadeh S, Dolman EM, Wieriks R, Sparidans RW, Hennink WE, Kok RJ. Anti-GD2 Immunoliposomes for Targeted Delivery of the Survivin Inhibitor Sepantronium Bromide (YM155) to Neuroblastoma Tumor Cells. Pharm Res. 2018;35(4):85. doi:10.1007/s11095-018-2373-x

44. Sallman DA, DeZern AE, Garcia-Manero G, et al. Eprenetapopt (APR-246) and Azacitidine in TP53-Mutant Myelodysplastic Syndromes. J Clin Oncol Off J Am Soc Clin Oncol. 2021;39(14):1584–1594. doi:10.1200/JCO.20.02341

45. McKinnell Z, Karel D, Tuerff D, Abrahim MS, Nassereddine S. Acute Myeloid Leukemia Following Myeloproliferative Neoplasms: A Review of What We Know, What We Do Not Know, and Emerging Treatment Strategies. J Hematol. 2023;11(6):197–209.

46. Lawrence L. Magrolimab/Azacitidine Combo Shows Efficacy for Untreated *TP53*-Mutant AML. Published online November 30, 2023. Accessed July 13, 2024. https://ashpublications.org/ashclinicalnews/news/7523/Magrolimab-Azacitidine-Combo-Shows-Efficacy-for

47. Sallman DA, Al Malki MM, Asch AS, et al. Magrolimab in Combination With Azacitidine in Patients With Higher-Risk Myelodysplastic Syndromes: Final Results of a Phase Ib Study. J Clin Oncol. 2023;41(15):2815–2826. doi:10.1200/JCO.22.01794

48. Thijssen R, Diepstraten ST, Moujalled D, et al. Intact TP-53 function is essential for sustaining durable responses to BH3-mimetic drugs in leukemias. Blood. 2021;137(20):2721–2735. doi:10.1182/blood.2020010167

49. Nishida Y, Ishizawa J, Ayoub E, et al. Enhanced TP53 reactivation disrupts MYC transcriptional program and overcomes venetoclax resistance in acute myeloid leukemias. Sci Adv. 2023;9(48):eadh1436. doi:10.1126/sciadv.adh1436

50. Velculescu VE, Madden SL, Zhang L, et al. Analysis of human transcriptomes. Nat Genet. 1999;23(4):387–388. doi:10.1038/70487

51. Hassin O, Nataraj NB, Shreberk-Shaked M, et al. Different hotspot p53 mutants exert distinct phenotypes and predict outcome of colorectal cancer patients. Nat Commun. 2022;13(1):2800. doi:10.1038/s41467-022-30481-7

