## Supplementary figures and images for "*BIRC5* dependency defines a targetable vulnerability in *TP53* mutant acute myeloid leukemia"

### Supplemental Figures

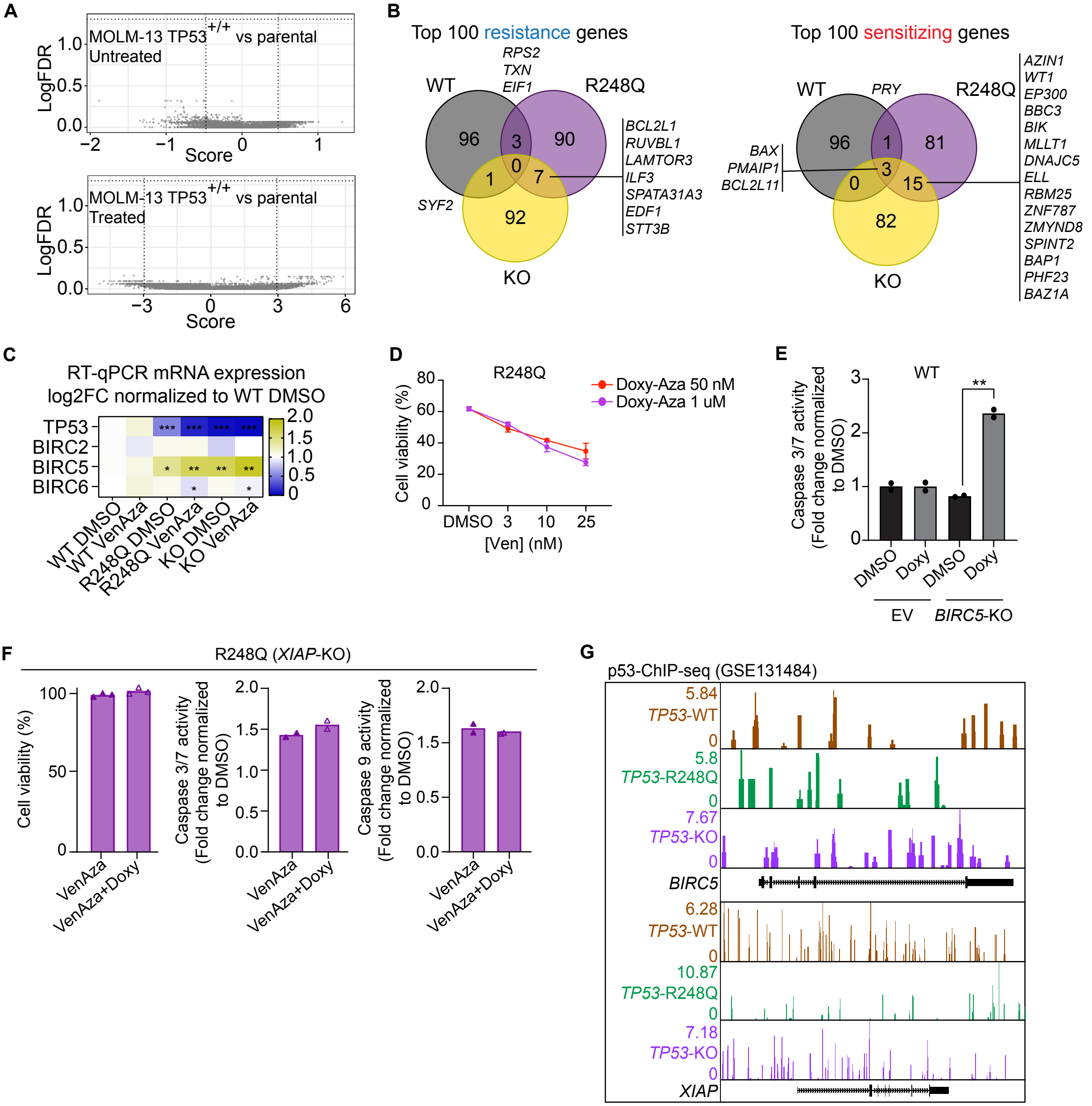

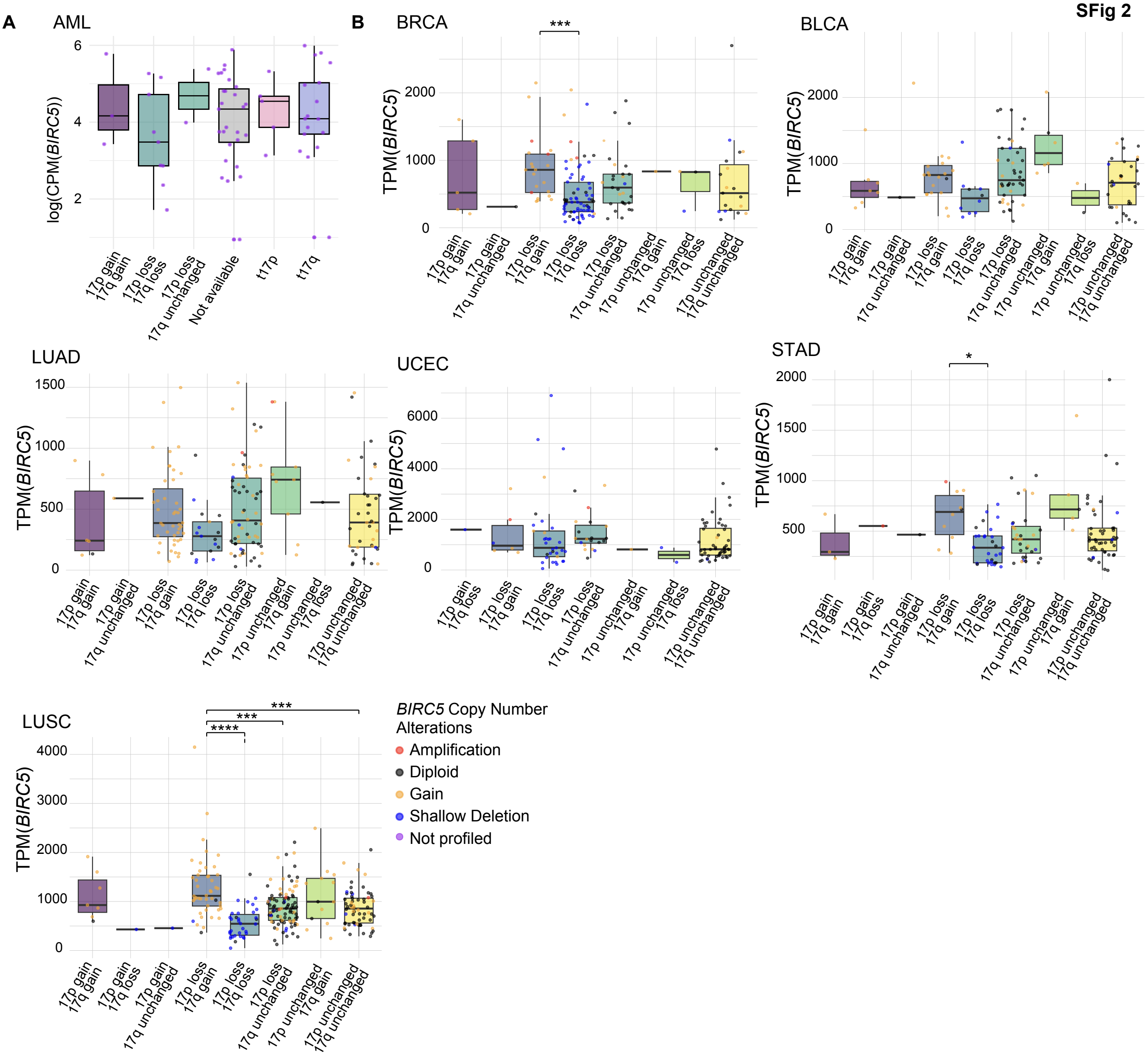

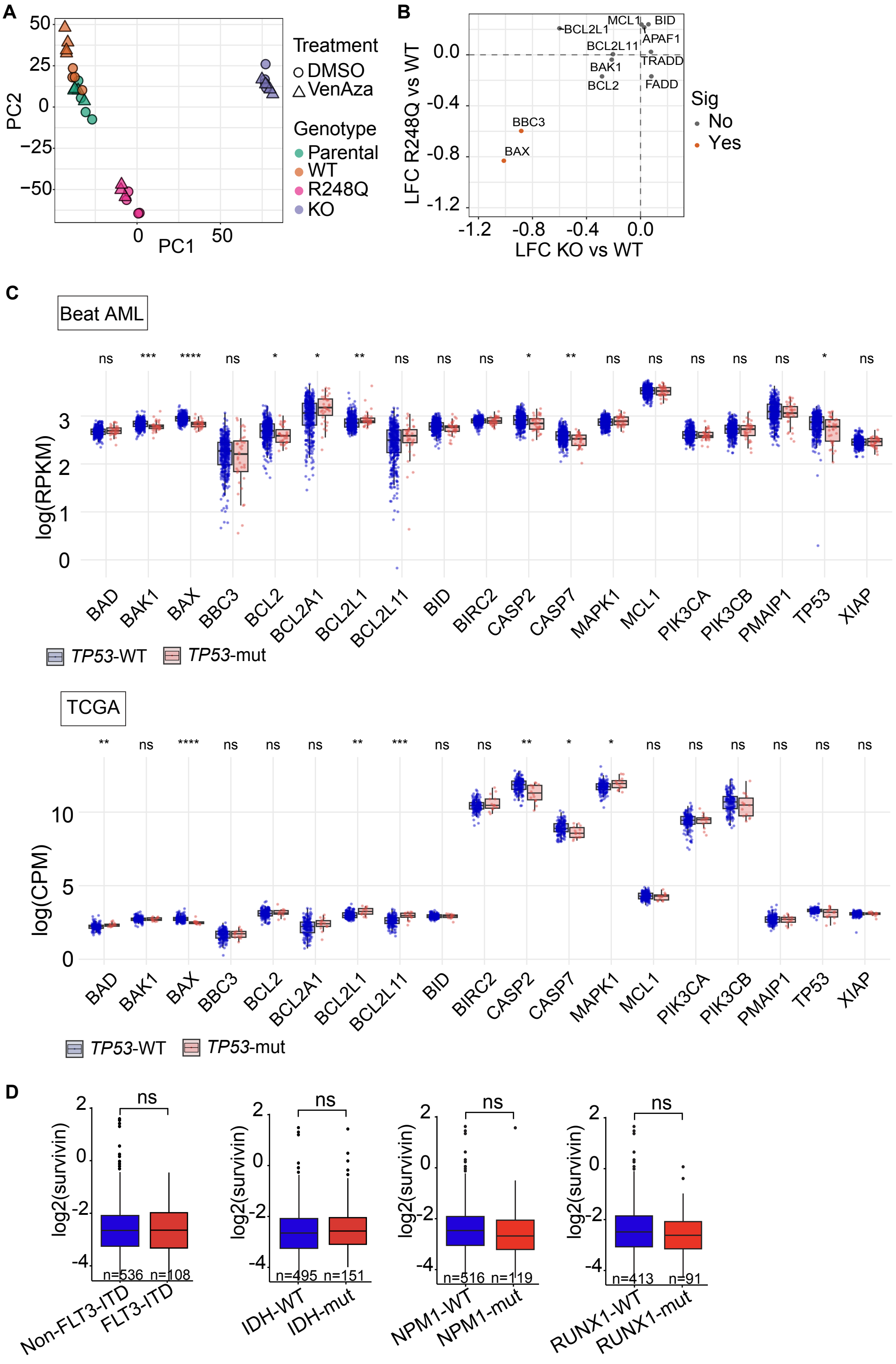

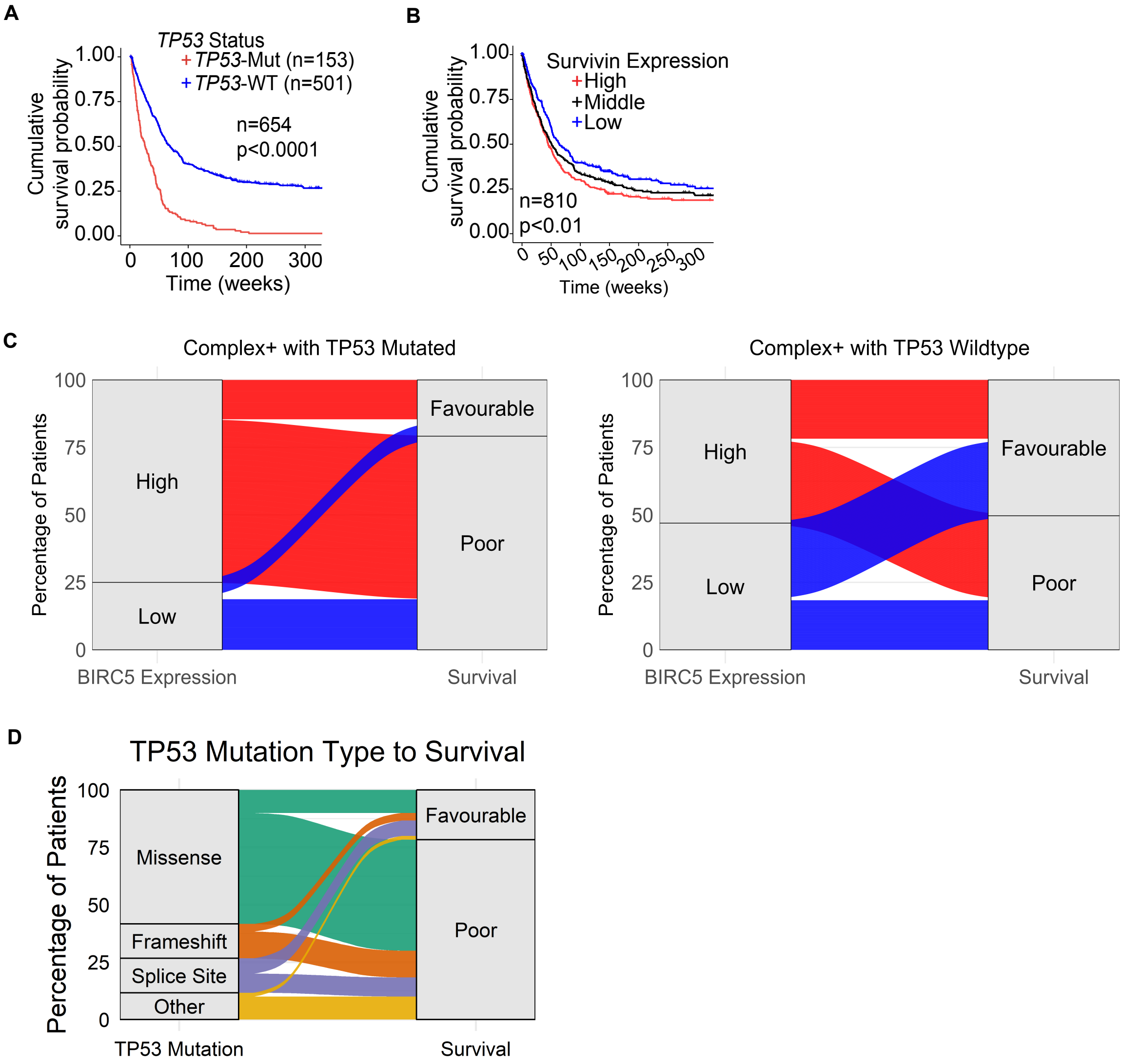

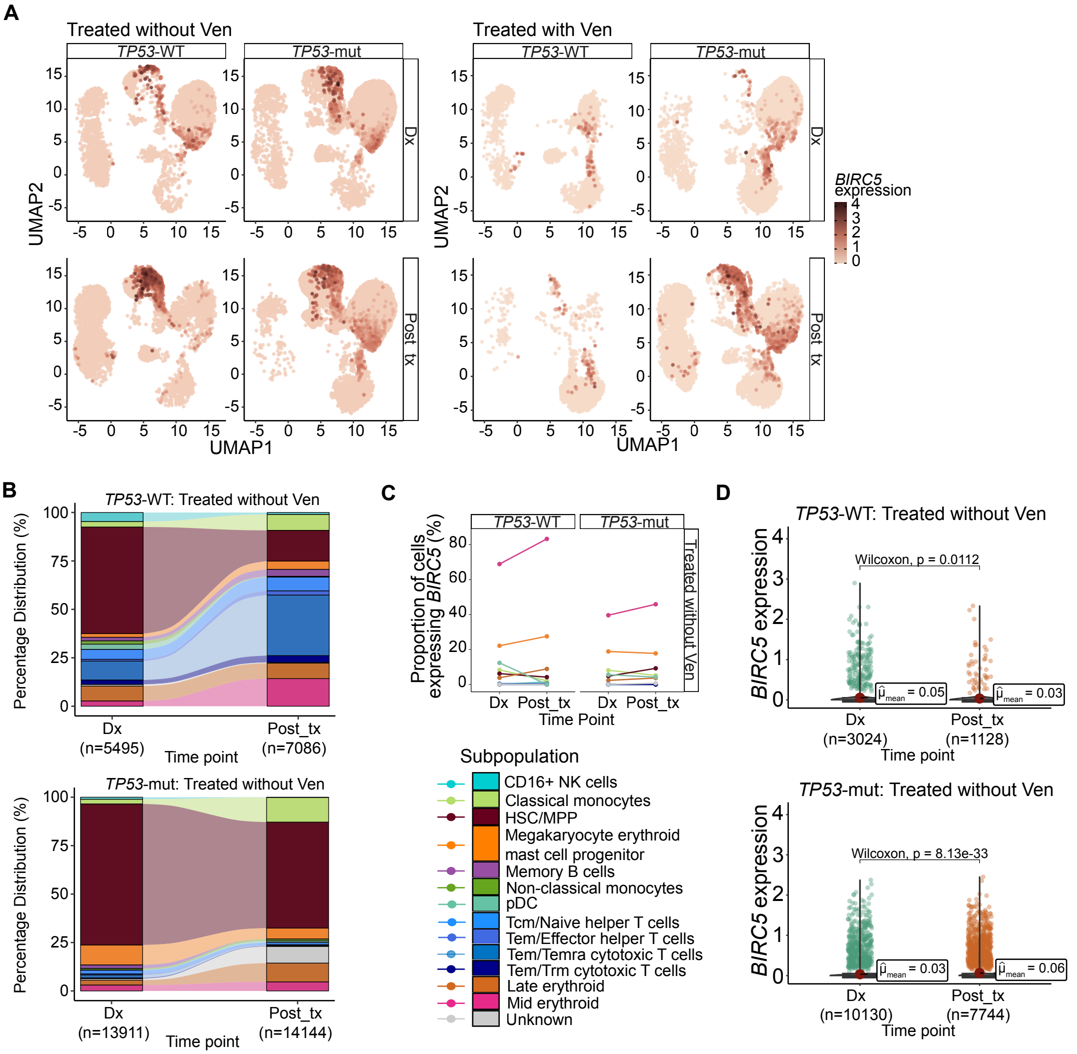

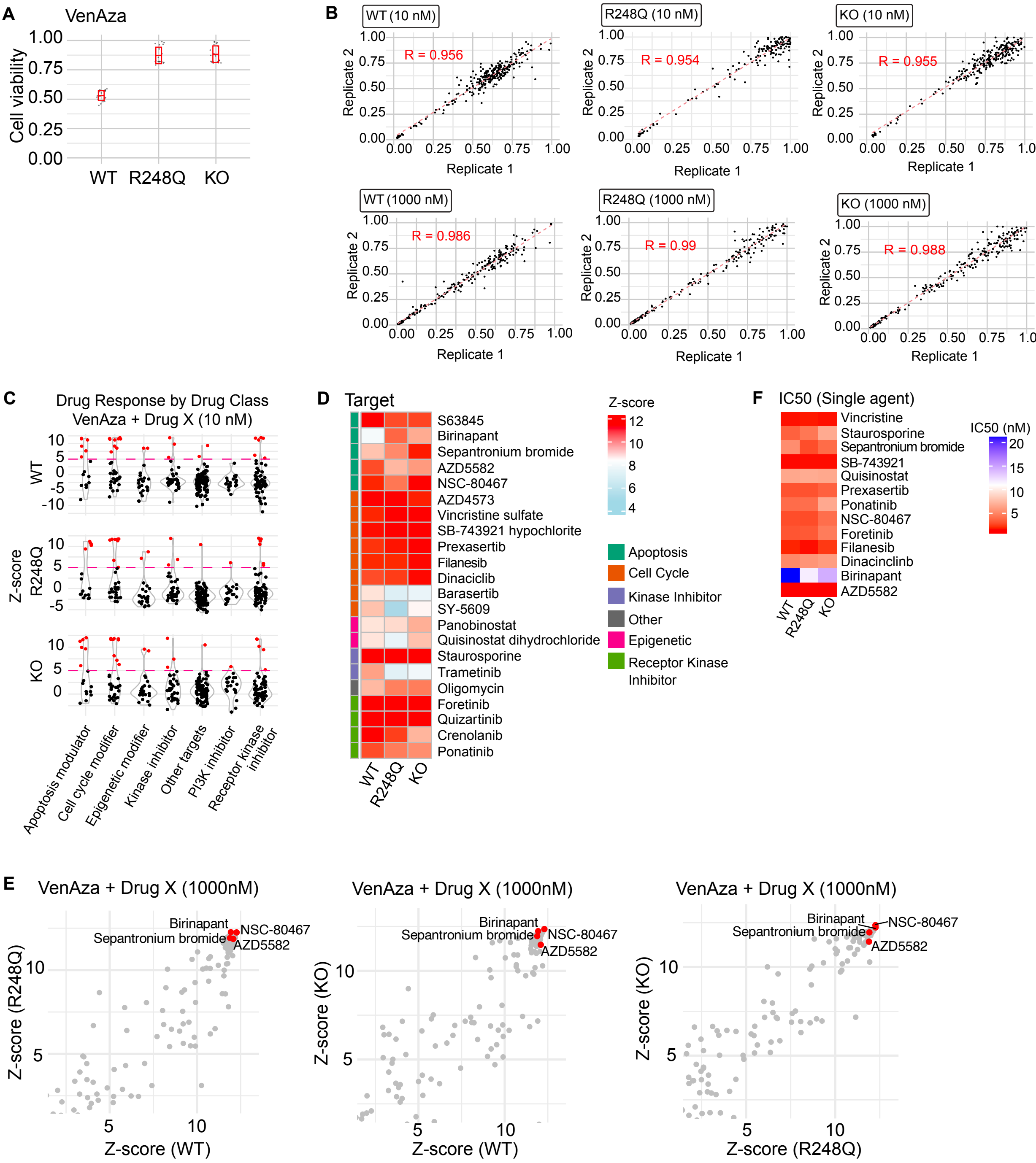

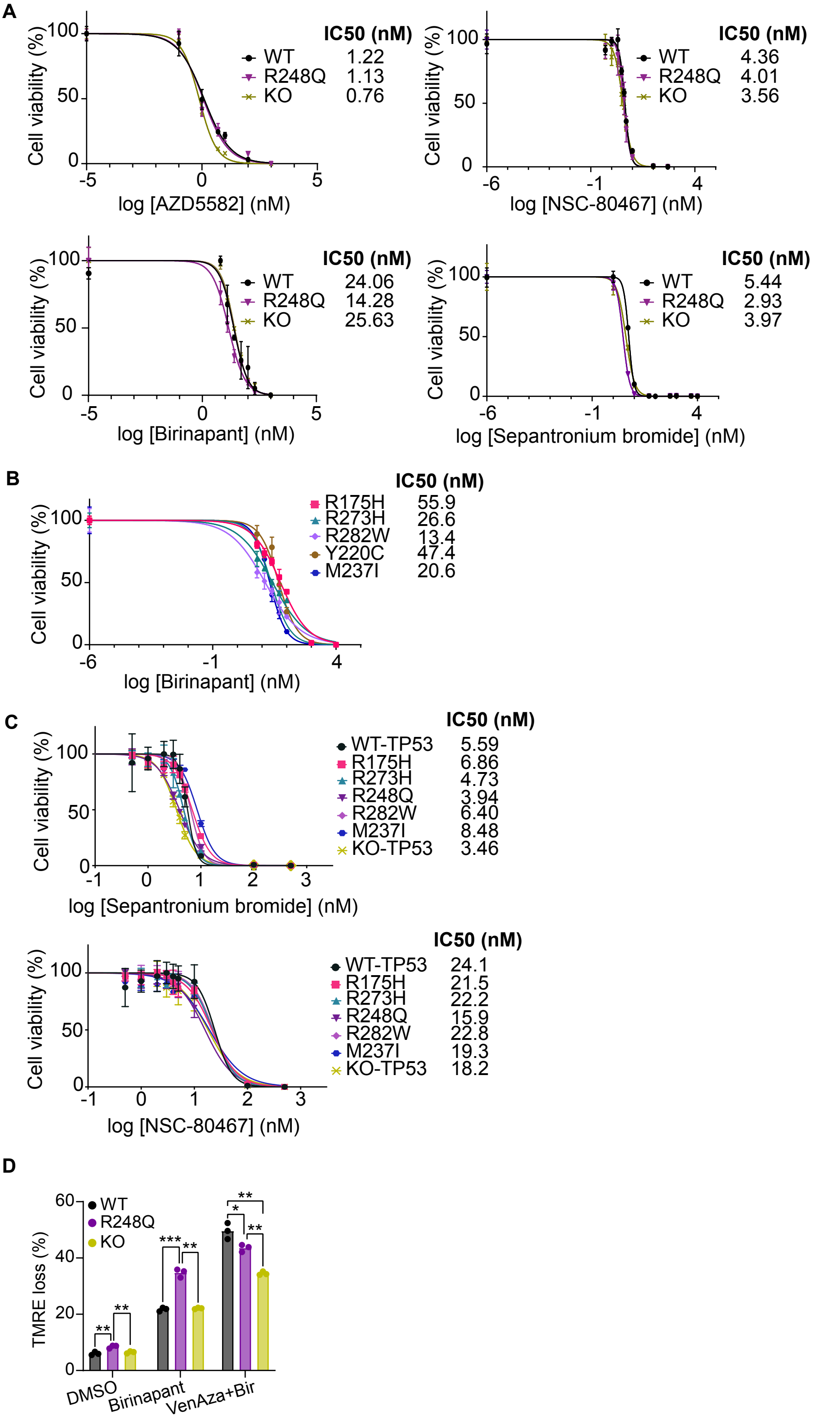

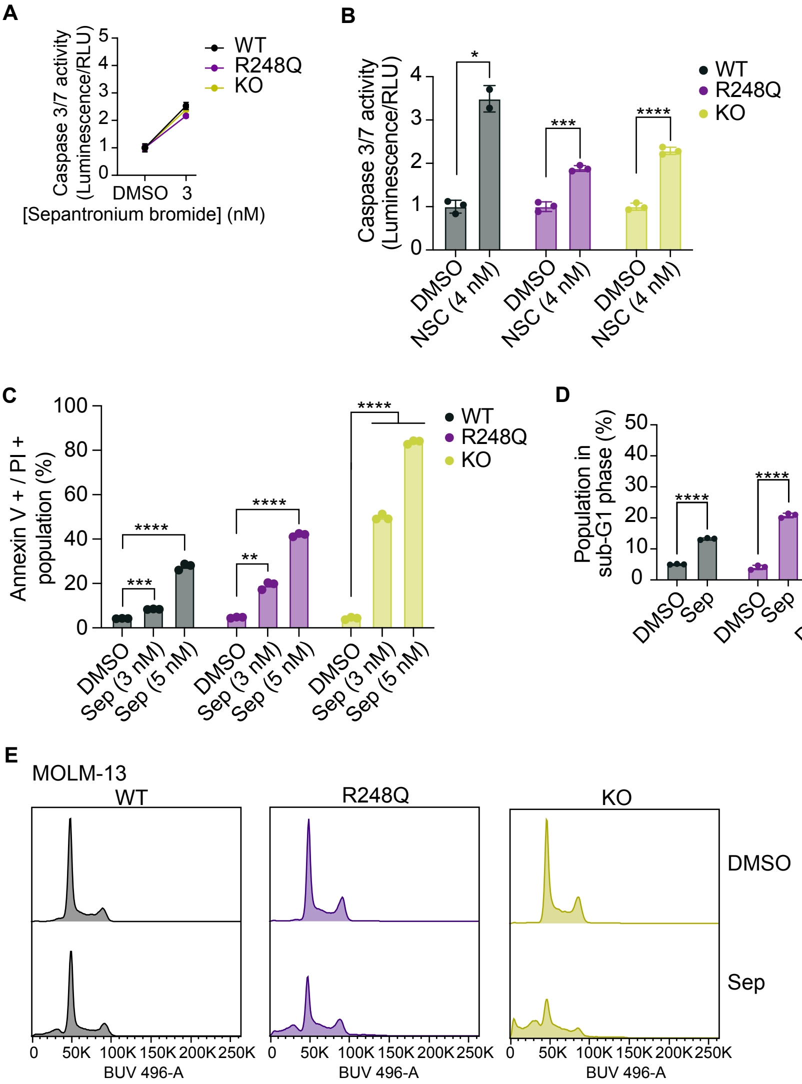
