## Supplemental Text for "*BIRC5* dependency defines a targetable vulnerability in *TP53* mutant acute myeloid leukemia"

### SUPPLEMENTAL METHODS

#### Animal studies

All animal studies were performed in accordance with protocols approved by the Institutional Animal Care and Use Committee (IACUC) guidelines. The DFAM-61345 PDX model is previously established by us and available from the Center for Patient-Derived Models at Dana-Farber Cancer Institute (https://www.pdxfinder.org/source/dfci-cpdm/). The PDX used is a quizartinib-resistant mouse with a gained *TP53* del17p. Female NSG (Jackson Laboratory) mice aged 6-8 weeks old were intravenously (IV) injected with $6.0\times10^5$ MOLM-13 cells or DFAM-61345.[22] To assess circulating leukemic burden, peripheral blood was collected and analyzed by flow cytometry, staining for human CD33 (clone WM53, BD Biosciences) and Zombie Yellow (BioLegend). Briefly, 10 µL of peripheral blood was lysed with 300 µL of ACK Lysing Buffer (Gibco). Data was acquired on a Cytoflex LX Flow Cytometer (Beckman Coulter) and analyzed using FlowJo softward. Drug formulations and administration schedules were as follows: Venetoclax (MedChemExpress) was prepared in a vehicle of 60% Phosal 50 PG, 30% PEG 400, and 10% EtOH and administered at 30 mg/kg orally (PO), five days per week. Azacitidine (MedChemExpress) was diluted in sterile 0.9 % NaCl and administered at 2.5 mg/kg intraperitoneally (IP), seven consecutive days during the first week of each 28-day cycle. Birinapant (MedChemExpress) was prepared in citrate buffer (pH 5.5) and administered at 25 mg/kg IP, three days per week. Sepantronium bromide was prepared in saline and administered at 1 mg/kg IP daily for ten days.

All animal studies were performed in accordance with approved IACUC guidelines at Dana-Farber cancer institute animal facility (IACUC protocol #14-038). DFAM-61786, DFAM-15354, DFAL-49600, DFAM-16835, DFAM-58159, DFAM-68555, and DFAM-61345 are available from the

Public Repository of Xenografts (PRoXe). Female NSG mice at 6-8 weeks of age (Jackson Labs) were injected with passage-2 0.6x106 human leukemia cells intravenously (I.V). Following transplant, mice were bled weekly, and treatment was initiated when circulating leukemia burden was >5% as assessed by flow cytometry staining for hCD45 (clone HI30, BD Biosciences) and hCD33 (clone WM53, BD Biosciences). All blood samples were lysed with ammonium chloride red-blood-cell buffer (Qiagen) prior to staining.

*Animal treatments*

Clinical grade venetoclax (Medchem express) was formulated in a mixture of 60% phosal 50 PG, 30% PEG 400, and 10% EtOH. Venetoclax was administered 100mg/kg per os (P.O.) five days per week. Quizartinib (Selleckchem) was formulated in 22% hydroxypropyl-β-cyclodextrin (Sigma-Aldrich, C0926), and administered 20mg/kg intraperitoneally (I.P.) five days per week. Birinapant (TargetMol Inc.) was formulated in citrate buffer (pH 5.5) and administered 5mg/kg I.P. three days per week. JQ-1 was formulated in 10% hydroxypropyl-β-cyclodextrin and administered 50mg/kg I.P. five days per week. JIB04 was formulated and dosed as previously described (2).

**Caspase luminescence assay**

A total of $1.0\times10^4$ MOLM-13 cells were seeded in 100 µL RPMI per well in 96-well plates and treated with the indicated agents. After 48 hours of incubation at 37°C, 30 µL of culture medium for each well was aliquoted to a 384-well plate. An equal volume (30 µL) of Caspase Glo reagent (Promega) was added to each well and incubated for 90 minutes at room temperature in the dark. Luminescence was measured using a Hidex Sense multi-plate reader.

**Cell culture**

AML cell lines were obtained as follows: MOLM-13 from DSMZ and MV-4-11 from ATCC. Isogenic MOLM-13-*TP53* cell lines: *TP53*$^{+/+}$ (wild-type), *TP53*$^{-/-}$ (knock-out), and six mutants (*TP53*$^{-/R248Q}$, *TP53*$^{-/Y220C}$, *TP53*$^{-/R82W}$, *TP53*$^{-/R173H}$, *TP53*$^{-/R275H}$, and *TP53*$^{-/M237I}$) were generously provided by Steffen Boettcher.[22] Lenti-X 293T cells were kindly provided by Nicholas Gascoigne. HCT-116 (*TP53*$^{+/+}$ and *TP53*$^{-/-}$) and CAL-51 (*TP53*$^{+/+}$ and *TP53*$^{-/-}$) cells were kindly provided by Chit Fang and Uri Ben-David, respectively. Isogenic MOLM-13 lines were cultured in RPMI-1640 (Gibco, 11875093). Lenti-X 293T, HCT-115, and CAL-51 cells were cultured in DMEM (Biowest, L0104). All media were supplemented with 10% fetal bovine serum (Gibco, 10437028) and 1% penicillin-streptomycin (Gibco, 15140122). Cells were cultured in a humidified incubator at 37°C with 5% $CO_2$.

**Cell cycle, apoptosis, and viability studies**

To assess cell cycle distribution, $5.0\times10^5$ cells were permeabilized and fixed using 70% ice-cold ethanol and stored at 4ºC. Cells were then washed with PBS, stained with 400 µL of DAPI containing 0.1% Triton-X, and incubated for 20 minutes at room temperature. Samples were analyzed by flow cytometry (LSRFortessa X-20) with excitation at an ultraviolet wavelength (496 nM). To measure apoptosis, $1.0\times10^4$ cells were seeded in triplicate in 96-well plates and treated with DMSO (vehicle control) or the indicated drug at 37°C for the designated duration. Cells were stained with Annexin V and propidium iodide solutions for ten minutes prior to acquisition on an Attune NxT Flow Cytometer. Data were analyzed using FlowJo software.

**CRISPR library screen**

A total of 1.0×10$^8$ MOLM-13 *TP53*$^{+/+}$, *TP53*$^{-/-}$, and *TP53*$^{-/R248Q}$ cells were transduced with the Human CRISPR Knockout Pooled Library (Brunello) (Addgene, #73179) at a multiplicity of infection (MOI) of 0.3 (#410).[24] Spin infection was performed at 2250 rpm for 30 minutes in the presence of polybrene at 10 µg/mL (MedChemExpress). Three days post-transduction, puromycin selection was applied for 72 hours. Eight days post-transduction, viable transduced cells were split into three treatment arms: untreated, DMSO (vehicle control), or venetoclax (3 nM) plus azacitidine (50 nM). Cells were maintained in culture under their respective conditions for 14 days and underwent four passages. Genomic DNA was extracted using phenol:chloroform:isoamyl alcohol (25:24:1) and purified with MaXtract High Density columns (QIAGEN). Library PCR amplification of sgRNAs was performed following Broad Institute protocols (PCR of sgRNAs for Illumina sequencing). Sequencing was conducted by on the NovaSeq X Plus platform (Azenta Life Sciences) at a sequencing depth of 500 reads per sgRNA. Reads counts of each sgRNA from the Brunello Human CRISPR Knockout Pooled Library[24] were quantified using the MAGeCK count module (v0.5.9.5)[25], as previously reported.[26] Gene-level enrichment was collapsed to the gene-level data and calculated with the MAGeCK test command using these parameters: *--gene-test-fdr-threshold 0.05*, *--adjust-method fdr,* and *--gene-lfc-method median*. Top-ranked gene dependencies visualized using MAGeCKFlute, as previously reported.[27]

**Cytotoxicity assay**

To assess cell viability, 1.0×10$^4$ cells were seeded in 96-well plates and treated with indicated agents at various concentrations. Cells were incubated for 72 hours at 37°C. Following incubation, CellTiter-Glo reagent (Promega) was added at a 4:1 ratio of cell culture medium to reagen/work solution, shaken vigorously for two minutes, and incubated for ten minutes at room temperature in the dark. Luminescence was measured using a Hidex Sense multi-plate reader. Absolute luminescence values (viability) were normalized to DMSO-treated controls to calculate percentage viability. IC50 values and dose-response curves were generated using a nonlinear regression analysis (log[inhibitor]) versus normalized response in GraphPad Prism.

**Data-independent acquisition (DIA) proteomics**

Frozen cell pellets were resuspended in 200 μL SDC lysis buffer and boiled at 95°C for ten minutes. Lysates were treated with benzonase (Merck) for 30 minutes, followed by overnight digestion at 37°C with trypsin (Promega) and LysC (Wako) at a 1:50 enzyme:protein ratio. Digestion was quenched by adding formic acid to a final concentration of 1%. Samples were centrifuged at maximum speed for 15 minutes at room temperature, and peptides were desalted using Sep-Pak C18 cc cartridges (Waters). Peptide concentrations were determined using the Pierce BCA Protein assay (ThermoFisher), and 1 μg of peptide was injected per run. Samples were analyzed using an Orbitrap Exploris 480 mass spectrometer (ThermoFisher) in Data Independent Acquisition (DIA) mode. Peptide separation was performed on a Vanquish Neo UHPLC system using a 110-minute gradient. Detailed LC-MS settings were as previously described.[28] Protein identification was conducted using DIA-NN (version 1.8.1[29]) in library-free mode with "reannotate" and "MBR" options enabled. The protein search was performed against the Uniprot human reference proteome (UP000005640). The DIA-NN output (pg file) was imported into R (version 4.3.2) for further analysis. The protein matrix was filtered to exclude contaminants and required a minimum of two unique peptides for proteins identification. Intensities were log2-transformed, and proteins with at least 70% valid values were retained. Missing values were imputed from a normal distribution (mean downshift = 1.8; width = 0.3 x SD). Differential protein expression analysis between experimental groups was performed using moderated two-sided t-tests implemented in the limma package (version 3.58.1[30]). All experiments were performed in quadruplicates, and p-values were adjusted for multiple testing using the Benjamini-Hochberg method.

**Drug library screen**

Technical duplicates of isogenic MOLM-13 *TP53*$^{+/+}$, *TP53*$^{-/R248Q}$, and *TP53*$^{-/-}$ cells were seeded in 96-well plates at $1.0\times10^4$ cells per well. Cells were treated with a combination of venetoclax (3 nM) and azacitidine (50 nM), along with individual agents from a curated 289-compound drug library at either 10 nM or 1000 nM concentrations. After 72 hours after incubation at 37°C, cell viability was measured using the CellTiter-Glo assay (Promega). Viability readouts were normalized across technical replicates, and DMSO-treated wells on either plate served as internal controls for baseline normalizations. Each drug was annotated by target pathway according to the Helsinki drug classification database. To assess differential response to VenAza, waterfall graph viability data (%) were converted into z-scores across the entire screen. A positive z-score indicated an enhanced sensitizing effect to VenAza, while a negative z-score indicated decreased sensitivity. Top candidate hits were defined as agents within the top 10% of z-scores for each cell line.

Equations: $Cell\ viability\ (\%) = \frac{CTG\ luminescence\ output\ of\ treated\ wells}{CTG\ luminescence\ output\ of\ DMSO\ control} * 100$

$$Z - score = \frac{(cell\ viability\ of\ VenAza\ -\ cell\ viability\ of\ triple\ combination)}{standard\ deviation\ of\ VenAza\ control}$$

$$Combination\ Index\ (CI) = \frac{D1}{(Dx)1} + \frac{D2}{(Dx)2}$$

D1 and D2 are doses of component 1 and component 2 in the combination, respectively. (Dx)1 and (Dx)2 are doses of component 1 and component 2 required to achieve a certain effect individually.[14] Synergistic index greater than 1 indicates synergy.

$$Synergistic\ Index\ (SI) = \frac{1}{Combination\ Index\ (CI)}$$

All drugs and responses can be found in Supplemental Table 1.

**Genetic editing in cell lines**

sgRNA sequences were cloned into lentiCRISPRv2 (Addgene, #52961) or FgH1tUTG (Addgene, #70183). For *TP53*$^{+/+}$ overexpression, lentiviral vectors pLV[Exp]-EGFP-CMV>hTP53 and pLV[Exp]-EGFP-CMV>ORF_Stuffer were custom-designed and packaged by VectorBuilder (VB230615-1583pby, VB900146-8110wvd). Lentiviral packaging was performed in Lenti-X 293T cells seeded at $1.8\times10^6$ cells per well. Cells were transfected with the envelope plasmid pVSV-G (415 ng), packaging plasmid psPAX2 (830 ng) and the corresponding transfer plasmid (1245 ng) using Lipofectamine™ 3000 (ThermoFisher, L3000015) and Opti-MEM (Gibco, 31985070) transfection reagents. After 24 hours, the culture media was replaced with fresh RPMI-1640. Lentiviral supernatants were harvested at 48 hours and 72 hours post-transfection and filtered through a 0.45 µM nitrocellulose filter (Sartorius). AML cells were transduced by centrifugation at 2500 rpm for 90 minutes at 32°C using 200 µL of virus-containing supernatant supplemented with 10 µg/mL polybrene (MedChemExpress). Antibiotic selection was initiated 48 hours post-infection using puromycin (2 µg/mL) or hygromycin (400 µg/ml), depending on the vector backbone. sgRNA sequences used in this study are listed in Supplemental Table 2.

**Patient cohorts**

Informed consent was obtained from all patients in compliance with the Declaration of Helsinki. RNA-seq data was from the Munich Leukemia Laboratory cohort (731 AML patient samples; *TP53*-wt, n = 668; *TP53*-mutant, n = 63). Proteomics profiling was conducted as previously described.[22] Nine *TP53* mutant primary tumors were obtained from the National University of Singapore, University of Miami, and MD Anderson Cancer Center (MDACC). Reverse Phase Protein Array was performed at the MDACC Core Facility following established protocols.[22–27] Peripheral blood and bone marrow samples were obtained from adult AML patients (n = 810) treated at MDACC. The study was approved by the MDACC IRB under protocols LAB01-473 and LAB05-0654.

**Reverse phase protein array**

Proteome profiling of over 200 proteins was performed as previously described.[37] Among a cohort of 810 patients, 413 samples (51.0%) were classified as *de novo* AML. Log2-normalized expression levels of survivin were compared between *TP53* wild-type samples (n = 501, 76.6%) and *TP53* mutant samples (n = 153, 23.4%).

**RT-qPCR and RNA-seq**

Total RNA was extracted using the RNeasy Mini Kit (QIAGEN) and treated with DNase I (QIAGEN) to remove genomic DNA contamination. For RT-qPCR, 1 µg of purified RNA was reverse-transcribed into cDNA using GoScript Reverse Transcriptase (Promega). Quantitative PCR was performed using Power SYBR Green PCR Master Mix (Applied Biosystems) on a CFX96 Touch Real-Time PCR Detection System (BioRad). Relative mRNA expression levels were calculated using the ΔΔ Ct method. Primer sequences are provided in Supplemental Table 3. For RNA sequencing, strand-specific libraries were constructed, and 100-bp paired-end sequencing was performed to a depth of 20 million reads per sample by Azenta Life Sciences. RNA-seq reads were aligned to the human reference genome GRCh38 with transcript annotations from GENCODE (version 45) using STAR aligner (version 2.7.11b.[38]). Gene-level raw counts were obtained with

STAR using the *--quantMode GeneCounts* option. Genes with fewer than ten total read counts across all samples were excluded. Differential gene expression analysis was estimated using DESeq2 (version 1.42.0.[39]).

**Single cell RNA-seq**

A total of 53 samples from 26 AML patients were collected and analyzed. Clinical characteristics, treatment regimens, and *TP53* mutation status for these samples are detailed in Supplemental Table 4. Single-cell RNA sequencing (scRNA-seq) was performed on a cohort of 56 bone marrow samples, including paired AML bone marrow samples and three normal bone marrow samples. Cells were dissociated and prepared for scRNA-seq using a standard protocol optimized for high viability and minimal cell loss. Single-cell libraries were generated using the Chromium Single Cell 3' Reagent Kits v3 (10x Genomics) following the manufacturer's instructions, and sequencing was performed on an Illumina platform. Data processing and quality control were carried out with Cell Ranger software (10x Genomics). Cells with at least 500 detected genes and less than 10% mitochondrial RNA content were retained for downstream analysis. Normalization, scaling, and identification of highly variable genes were performed using the Scanpy package (version 1.7.2). Cells from all samples were clustered via the Leiden algorithm, and cluster annotation was conducted with CellTypeST tool, leveraging known marker genes to accurately classify hematopoietic and leukemic subpopulations. The relative abundance of different cell populations – including hematopoietic stem and progenitor cells (HSPCs), erythroid cells, and various myeloid lineages – was quantified and compared across samples. For *BIRC5* expression analysis, paired samples (n = 31) from 15 *TP53* mutant and paired samples (n=14) from seven *TP53* wild-type AML patients were selected to ensure consistent *TP53* mutation status at diagnosis and post-treatment timepoints. Normalized *BIRC5* expression was used for all comparisons by *TP53* mutation status and treatment type. Single cells were labelled as *BIRC5*-expressing if their normalized *BIRC5* expression was greater than zero. Statistical comparisons were performed using the Wilcoxon rank-sum test.

**TMRE staining**

Cells were seeded at a density of $1.0 \times 10^6$ cells per mL of culture media. Cells were stained with TMRE (10 nM, Invitrogen) and incubated at 37°C with 5% $CO_2$ for 30 minutes. After staining, cells were pelleted, resuspended in fresh media, and treated as indicated. Following treatment, cells were incubated for an additional 24 hours before being stained with DAPI prior to analysis by flow cytometry.

**Western blot**

Cells were lysed in NP-40 lysis buffer (150 mM NaCl, 1% NP-40, 50 mM Tris-Cl pH 8.0; Cell Signaling Technology) supplemented with protease inhibitor for 45 minutes at 4°C, following the manufacturer's protocol. Mitochondria were isolated using the Mitochondria Isolation Kit (ThermoFisher), and protein concentration was determined using the Pierce BCA Protein Assay Kit Kit (ThermoFisher). For Western Blot: Proteins were denatured in Laemmli Sample Buffer (Bio-Rad) containing 10% β-mercaptoethanol (Sigma-Aldrich), separated by SDS-PAGE, and transferred onto nitrocellulose membranes. Membranes were blocked for one hour in 5% milk (Bio-Rad) dissolved in TBST and incubated overnight with primary antibodies diluted in 5% milk. Densitometric quantification of western blot bands was performed using FlowJo software. The list of antibodies used is detailed in Supplemental Table 5.

### SUPPLEMENTAL TABLES

**Supplemental Table 1.** Agents from HT drug screening.

**Supplemental Table 2.** sgRNA sequences used in CRISPR-KO experiments.

**Supplemental Table 3.** Primer sequences used for RT-qPCR

**Supplemental Table 4.** Clinical details, type of treatment received, and *TP53* mutation status of samples from single-cell RNA-seq.

**Supplemental Table 5.** List of antibodies used.

### SUPPLEMENTAL FIGURE LEGENDS

**Supplemental Figure 1.**

**(A)** Functional dependencies in a single $TP53^{+/+}$ clone closely match those of parental MOLM-13 cells. **(B)** Venn diagram of top 100 sensitizing and resistance genes across all genotypes. **(C)** Heatmap representing mRNA expression of IAP genes obtained from RNA-seq of isogenic MOLM-13 cells normalized to $TP53^{+/+}$ cells treated with DMSO. **(D)** *BIRC5*-KO shows sensitivity to azacitidine and venetoclax. **(E)** *BIRC5*-KO in $TP53^{+/+}$ cells induces restoration of caspase-3/7 activity. All data was normalized to DMSO (no-doxycycline)-treated cells. **(F)** Cell viability, caspase-3/7, and caspase-9 activity in *XIAP*-KO $TP53^{-/R248Q}$ cells treated with VenAza with or without doxycycline (10 ng) for 72 hours. All data was normalized to DMSO (no-Doxy)-treated cells. **(G)** ChIP-seq gene tracks (GSE131484) depict lack of p53 binding at *BIRC5* and *XIAP* loci in $TP53^{+/+}$, $TP53^{-/R248Q}$, and $TP53^{-/-}$ cells. For all figures unless otherwise specified, treatment with VenAza: venetoclax 3 nM, azacitidine 50 nM. ns = no significance, *p<0.05, **p<0.01, ***p<0.001, ****p<0.0001.

**Supplemental Figure 2.**

**(A)** *BIRC5* expression levels of patients with *TP53* mutations across cancers from the TCGA dataset, stratified according to chromosome 17 CNV. Analysis of samples using one-tailed Wilcoxon signed-rank test with Bonferroni correction shows that mean expression of *BIRC5* in patients with 17p loss and 17q gain may be significantly greater than that of other CNVs. Analysis via Wilcoxon test. ns = no significance, *p<0.05, **p<0.01, ***p<0.001, ****p<0.0001.

**Supplemental Figure 3.**

**(A)** PCA plot of proteomics data from isogenic MOLM-13 cells. Each shape represents a treatment replicate (circle = DMSO, triangle = VenAza) (n = 4), colored by genotype (green = parental, orange = $TP53^{+/+}$, pink = $TP53^{-/R248Q}$, purple = $TP53^{-/-}$). One replicate ($TP53^{-/R248Q}$, VenAza) was removed due to significant difference in read counts compared to its counterparts. **(B)** Integration of common genes from RNAseq and proteins from proteomics analysis (n = 106) for $TP53^{-/R248Q}$ versus $TP53^{+/+}$ and $TP53^{-/-}$ versus $TP53^{+/+}$ cells treated with VenAza. **(C)** mRNA expression levels of apoptosis genes obtained from RNA-seq of individual AML patient samples from Beat AML and TCGA datasets, stratified according to *TP53* status. Analysis via Wilcoxon test. **(D)** RPPA protein expression of survivin in AML patients from MD Anderson Cancer Center by gene status showing *FLT3, IDH, NPM1,* and *RUNX1* mutations does not affect survivin levels. Analysis via Wilcoxon test. For all figures unless otherwise specified, treatment with VenAza: venetoclax 3 nM, azacitidine 50 nM. ns = no significance, *p<0.05, **p<0.01, ***p<0.001, ****p<0.0001.

**Supplemental Figure 4.**

**(A-B)** Kaplan-Meier survival plot and analysis reveals that cumulative survival probability for AML groups from the MD Anderson Cancer Center cohort is stratified according to **(A)** *TP53* status and **(B)** survivin levels. Analysis via Wilcoxon test. ns = no significance, *p<0.05, **p<0.01,

***p<0.001, ****p<0.0001. **(C)** Sankey diagram showing the association between *BIRC5* expression levels (High vs Low) and survival outcomes (Favorable vs Poor) in patients with complex karyotypes and mutant *TP53* (n = 48) vs wild-type *TP53* (n = 147). Flow widths represent the proportion of patients within each category. **(D)** *TP53* mutant patients (n = 60) stratified by type of mutation (missense (n = 35), frameshift (n = 9), splice (n = 9), other (nonsense and insertions/deletions; n = 7), and wild-type) shows that missense and frameshift mutants have worse overall survival compared to other types (p<0.05). For figures C-D, analysis via Wilcoxon rank-sum test. *BIRC5* expression stratified into "high" or "low" on the basis of median expression levels within patients with *TP53* mutations. Survival status was "poor" (overall survival ≤ 360 days) or "favorable" (> 360 days), with an observed cohort median of 360.5 days used as the cut-off. Thickness of flows indicates the number of patients in each survival–mutation category.

**Supplemental Figure 5.**

**(A)** UMAP plot visualizes proportion of cells treated without venetoclax expressing *BIRC5* (defined as *BIRC5* gene expression > 0) in in various immune cell subpopulations between two timepoints (diagnosis (Dx) and post-treatment (Post-Tx)) by *TP53* mutation status. **(B)** Sankey diagram illustrates the shift in percentage distribution of various cell subpopulations treated without venetoclax between two time points (diagnosis (Dx) and post-treatment (Post-Tx)) by *TP53* mutation status. For *TP53*-WT (n = 6 patients), single cells are n = 3153 (Dx) and n = 1535 (Post-Tx). For *TP53*-Mut (n = 9 patients), single cells are n = 10314 (Dx) and n = 7819 (Post-Tx). Wilcoxon signed rank test. **(C)** Proportion of cells treated without venetoclax expressing *BIRC5* (defined as *BIRC5* gene expression > 0) in various cell subpopulations between two timepoints (diagnosis, Dx and post-treatment, Post-Tx) by *TP53* mutation status. **(D)** Distribution of single cells based on their *BIRC5* gene expression profiles for *TP53*-WT and *TP53*-Mut cells without combination of venetoclax For *TP53*-WT n = 6 patients), single cells are n = 5652 (Dx) and n = 9015 (Post-Tx). For *TP53*-Mut (n = 9 patients), single cells are n = 17414 (Dx) and n = 19519 (Post-Tx). ns = no significance, *p<0.05, **p<0.01, ***p<0.001, ****p<0.0001.

**Supplemental Figure 6.**

**(A)** Cell viability of MOLM-13 isogenic $TP53^{+/+}$, $TP53^{-/R248Q}$, and $TP53^{-/-}$ cells after VenAza treatment for 72 hours was compared across 20 plates. **(B)** Pearson correlation analysis of technical replicates in drug screens. **(C)** Z-score correlation scatterplots comparing z-scores of agents at 10 nM by drug class, with comparisons between $TP53^{-/R248Q}$ vs $TP53^{+/+}$, and $TP53^{-/-}$ vs $TP53^{+/+}$. Z-score cut-off > 2 across all genotypes. **(D)** Heatmap of z-score drug sensitivities for clinically relevant agents at 10 nM. **(E)** Z-score correlation between $TP53^{-/R248Q}$ vs $TP53^{+/+}$, $TP53^{-/-}$ vs $TP53^{+/+}$, and $TP53^{-/R248Q}$ vs $TP53^{-/-}$ identifies agents specific to *TP53* status at 1000 nM. **(F)** Single agent (monotherapy) IC50 values of selected agents determined in $TP53^{+/+}$, $TP53^{-/R248Q}$, and $TP53^{-/-}$ cells. For all figures unless otherwise specified, treatment with VenAza: venetoclax 3 nM, azacitidine 50 nM. Experiments were repeated twice. One biological replicate is shown with three technical replicates.

**Supplemental Figure 7.**

**(A)** Cell viability plots for $TP53^{+/+}$, $TP53^{-/R248Q}$, and $TP53^{-/-}$ cells treated with AZD5582, birinapant, NSC-80467, or sepantronium bromide for 72 hours. **(B-C)** $TP53^{-/mut}$ cells treated with **(B)** birinapant and **(C)** sepantronium bromide or NSC-80467 for 72 hours exhibit decreased viability. **(D)** TMRE loss for $TP53^{+/+}$, $TP53^{-/R248Q}$, $TP53^{-/-}$ cells treated with DMSO, birinapant (20 nM), or VenAza-Bir for 24 hours. For all figures unless otherwise specified, treatment with VenAza:

venetoclax 3 nM, azacitidine 50 nM. ns = no significance, *$p<0.05$, **$p<0.01$, ***$p<0.001$, ****$p<0.0001$.

**Supplemental Figure 8.**

**(A-B)** Caspase-3/7 activity increases in *TP53*$^{+/+}$, *TP53*$^{-/R248Q}$, and *TP53*$^{-/-}$ cells treated with **(A)** sepantronium bromide or **(B)** NSC-80467 (4 nM) or 72 hours. **(C)** Annexin V-APC and PI assay shows increased cell death populations in *TP53*$^{+/+}$, *TP53*$^{-/R248Q}$, and *TP53*$^{-/-}$ cells treated with sepantronium bromide (3 nM, 5 nM) for 72 hours. **(D-E)** MOLM-13 *TP53*$^{+/+}$, *TP53*$^{-/\mathrm{R248Q}}$, and *TP53*$^{-/-}$ cells treated with DMSO or sepantronium bromide for 48 hours, **(D)** percentage of cell populations in sub-G1 phase, and **(E)** cell cycle profiles. For all figures unless otherwise specified, treatment with sepantronium bromide 3 nM. ns = no significance, *$p<0.05$, **$p<0.01$, ***$p<0.001$, ****$p<0.0001$.

**Supplemental Figure 9.**

**(A)** *TP53*$^{+/+}$ mice post-transplant treated with birinapant monotherapy derive no therapeutic benefits compared to untreated mice. **(B)** Survival percentage of *TP53*$^{+/+}$ mice treated with VenAza and *TP53*$^{-/\mathrm{R248Q}}$ mice treated with VenAza-Bir as proof-of-concept for triplet therapy. Treatment started five days post engraftment. **(C)** Weights of mice post-treatment with VenAza or VenAza-Bir shows overall good tolerability of agents. **(D)** Representative histology of Haemotoxylin and Eosin staining of *TP53*$^{-/\mathrm{R248Q}}$ mice exposed to VenAza-Bir for six weeks for lung, spleen, liver, bone marrow, heart, and kidney. For *in vivo* dosing: venetoclax 30 mg/kg PO, azacitidine 2.5 mg/kg IP, birinapant 25 mg/kg IP, sepantronium bromide 1 mg/kg IP.

**Supplemental Figure 10.**

**(A)** *TP53*$^{+/+}$ and *TP53*$^{-/-}$ CAL-51 and HCT-116 cells treated with doxorubicin or 5-FU, respectively. Cell viability is greater in *TP53*$^{-/-}$ cells. **(B-C)** Cell viability plots for *TP53*$^{+/+}$ and *TP53*$^{-/-}$ CAL-51 and HCT-116 cells treated for 72 hours with **(B)** NSC-80467 and **(C)** birinapant alone (+DMSO, 1 μM, 5 μM, 10 μM, and 50 μM) or with chemotherapy (+5-FU or +Dox). For all figures unless otherwise specified, treatment with doxorubicin 50 nM and 5-FU 50 μM. ns = no significance, *p<0.05, **p<0.01, ***p<0.001, ****p<0.0001.
